# *Dmc1* is a candidate for temperature tolerance during wheat meiosis

**DOI:** 10.1101/759597

**Authors:** Tracie Draeger, Azahara Martin, Abdul Kader Alabdullah, Ali Pendle, María-Dolores Rey, Peter Shaw, Graham Moore

## Abstract

We have assessed the effects of high and low temperatures on meiotic chromosome synapsis and crossover formation in the hexaploid wheat (*Triticum aestivum* L.) variety ‘Chinese Spring’. At low temperatures, asynapsis and chromosome univalence have been observed before in Chinese Spring lines lacking the long arm of chromosome 5D (5DL), which led to the proposal that 5DL carries a gene (*Ltp1*) that stabilises wheat chromosome pairing at low temperatures. In the current study, Chinese Spring wild type and 5DL interstitial deletion mutant plants were exposed to low (13°C) or high (30°C) temperatures in controlled environment rooms during a period from premeiotic interphase to early meiosis I. A 5DL deletion mutant was identified whose meiotic chromosomes exhibit extremely high levels of asynapsis and chromosome univalence at metaphase I after seven days at 13°C. This suggests that the mutant, which we name *ttmei1* (*t*emperature *t*olerance in *mei*osis 1) has a deletion of a gene that, like *Ltp1*, normally stabilises chromosome pairing at low temperatures. Immunolocalisation of the meiotic proteins ASY1 and ZYP1 on *ttmei1* mutants showed that low temperature results in a failure to complete synapsis at pachytene. After 24 hours at 30°C, *ttmei1* mutants exhibited a reduced number of crossovers and increased univalence, but to a lesser extent than at 13°C. KASP genotyping revealed that *ttmei1* has a 4 Mb deletion in 5DL. Of 41 genes within this deletion region, the strongest candidate for the stabilisation of chromosome pairing at low (and possibly high) temperatures is the meiotic recombination gene *Dmc1*.

**Key message:** The meiotic recombination gene *Dmc1* on wheat chromosome 5D has been identified as a candidate for the maintenance of normal chromosome synapsis and crossover at low and possibly high temperatures.

## Introduction

In plants, male reproductive development is highly sensitive to adverse environmental conditions including high and low temperatures (De Storme and Geelen 2014). In wheat, the reproductive phase is more sensitive to high temperatures than the vegetative phase (Fischer and Maurer 1976). This is also the case for low temperatures, with meiosis I identified as the most sensitive stage (Thakur et al. 2010). High temperatures during reproductive development can have a negative impact on grain yield (Fischer and Maurer 1976; Fischer 1985; Wardlaw et al. 1989), and even short periods of a moderately high temperature (20-24 hours at 30°C) during meiosis can reduce grain number (Saini and Aspinall 1982; Draeger and Moore 2017). Grain yield is also reduced when low temperatures occur during the booting stage (Ji et al. 2017), which broadly corresponds to meiosis (Barber et al. 2015).

In cereal crops such as wheat, where grains are an important yield factor, stress-induced male sterility generally has a negative effect on crop yield and performance (De Storme and Geelen 2014). In the UK and France in the mid-1980s, cold and wet weather during floral development in the wheat variety Moulin caused significant sterility, resulting in a reduction in grain yield of more than 70% (Law 1999). The dramatic falls in yield were thought to be a result of reduced fertility caused by the occurrence of low temperatures at meiosis. It was suggested that Moulin might carry specific alleles from diverse sources that made the wheat more sensitive to cold weather during meiosis.

Meiosis is essential for gamete formation in sexually reproducing organisms. In early meiosis, homologous chromosomes align next to each other as pairs and then, during synapsis, the pairs of homologs become linked tightly with each other through the polymerization of a protein structure called the synaptonemal complex (SC), which assembles between the paired chromosomes (Page and Hawley, 2004). The SC has a ladder-like structure consisting of two chromosome axes and a central region. Its assembly can be tracked by immunolocalisation of the meiotic proteins ASY1, which interacts with the chromosome axes (Boden et al. 2009), and ZYP1, which is associated with the central region (Higgins et al. 2005; Khoo et al. 2012). During zygotene the axes of the two homologs begin to become connected by the assembly of the central region between them. The axes are now called lateral elements. At pachytene the central structure links the homologs along their entire lengths and synapsis is completed. SC assembly is a highly temperature-sensitive process (Bilgir et al. 2013).

The SC is thought to provide the structural framework for meiotic recombination to take place. During recombination, at least one crossover (CO) must form between each pair of homologs, to forge a physical connection between the chromosomes. These physical connections can be seen cytologically and are called chiasmata. COs enable genetic information to be exchanged between chromosomes and are also needed for accurate chromosome segregation and balanced gametes in the daughter cells. Once COs have fully formed (at pachytene), the SC is disassembled (at diplotene), at which point the homologs are only connected via their chiasmata. They remain connected until the chromosomes segregate at anaphase I.

In bread wheat, which is an allopolyploid with three homeologous (related) diploid genomes (AABBDD), COs only form between homologs and not between homeologs, ensuring that the three genomes behave as diploids during meiosis. Normally, at metaphase I, only ring bivalents (with two chiasmata) and occasional rod bivalents (one chiasma) are present. At metaphase I the bivalents align on the equatorial plate before segregating at anaphase I.

Correct pairing and segregation of homologs is vital for maintaining the stability and fertility of the genome. Errors in meiosis can lead to aneuploidy or infertility. High and low temperatures can induce a variety of meiotic aberrations in plants including changes in the frequency of chiasma formation (CO frequency) (Elliott 1955; Dowrick 1957; Bayliss and Riley 1972a; Higgins et al. 2012). Reduction in chiasma formation at extremes of temperature is linked to disruption of chromosome synapsis. This can result in unpaired univalent chromosomes that segregate randomly during meiosis I or are lost completely (Bomblies et al. 2015). Temperature-associated synapsis failure has been reported in plants at both low and high temperatures, but the temperature at which meiosis fails varies in different species (reviewed in Bomblies et al. 2015). These temperature thresholds can also vary within a species where there are genotypic differences (Riley et al. 1966).

Chinese Spring is one of the more heat-sensitive wheat cultivars (Qin et al. 2008) and has been widely used to study the frequency of chiasma formation at low and high temperatures because different Chinese Spring genotypes respond differently to temperature extremes. At low temperatures, a reduction in chiasma frequency and an increase in chromosome univalence has been observed in Chinese Spring nullisomic 5D-tetrasomic 5B (N5DT5B) plants, which lack chromosome 5D and have two extra copies of 5B (Riley 1966; Bayliss and Riley 1972a). Wheat has an optimum temperature range of around 17-23°C over the course of a growing season (Porter and Gawith 1999). In N5DT5B plants, the frequency of chiasma formation is progressively reduced as the temperature rises above or falls below the optimum range (Bayliss and Riley 1972a). At 15°C, chiasma frequency is greatly reduced in N5DT5B plants (Riley 1966), and they exhibit pronounced chromosome pairing failure at 12°C, which leads to complete male sterility (Hayter and Riley 1967). The reduced chiasma frequencies seen in N5DT5B plants at low temperatures occur because the chromosomes fail to pair during zygotene, though the temperature sensitive phase was found to be during premeiotic interphase, prior to DNA synthesis (Bayliss and Riley 1972b).

Chromosome pairing failure in N5DT5B is not a result of the extra dosage of chromosome 5B (Riley et al. 1966), so it was inferred that there must be a gene on chromosome 5D that stabilises chromosome pairing at low temperatures. This gene was named *Ltp* (*Low temperature pairing*) by Hayter and Riley (1967). *Ltp* was further defined to the long arm of chromosome 5D (Hayter 1969) and later renamed *Ltp1* (Queiroz et al. 1991), but its exact location has never been mapped. It has been suggested that chiasma frequency also falls progressively in N5DT5B plants at temperatures of 30°C and above (Bayliss and Riley 1972a), indicating that chromosome 5D may also be associated with high temperature tolerance. However, this suggestion was based on the scoring of only a few cells because exposure of the plants to high temperatures for three days made the chromosomes too sticky to score accurately. Grain number is also reduced much more in N5DT5B plants than in the wild type after exposure to 30°C during premeiosis and leptotene (Draeger and Moore 2017).

The main aims of this study were to define *Ltp1* to a small enough region on 5DL to enable the identification of a candidate gene for the low temperature pairing phenotype, and to determine whether tolerance to both high and low temperatures could be controlled by the same locus. The strategy for mapping *Ltp1* was similar to the one used to map the *Ph1* locus (Roberts et al. 1999; Griffiths et al. 2006) a major locus that promotes homolog synapsis and regulates crossover formation in wheat (Martín et al. 2014, 2017). This had involved screening for gamma irradiation induced deletions on specific chromosomes in large populations of wheat. The work on *Ph1* established the dose of gamma irradiation likely to give a good rate of deletion discovery. This strategy was applied in the current study to identify deletions of chromosome 5DL using chromosome specific markers. Recent development of resources such as the Chinese Spring IWGSC RefSeq v1.0 genome assembly (International Wheat Genome Sequencing Consortium [IWGSC] 2018), the Wheat 820K Axiom® Breeders’ Array probe set (Winfield et al. 2016) and the Ensembl Plants database (Bolser et al. 2016) facilitated the processes of mapping and candidate gene identification.

## Materials and methods

### Plant materials

Deletion lines were generated in bread wheat (*Triticum aestivum* L., 2n = 6x = 42) var. ‘Chinese Spring’ by gamma irradiation of 1000 seeds at the International Atomic Energy Agency, Vienna (500 seeds irradiated at 250Gy and 500 at 300Gy). M_1_ deletion lines were self-fertilised. M_2_ lines were genotyped using KASP (Kompetitive allele specific PCR) to identify plants with interstitial deletions of 5DL. These deletion mutants were exposed to 13°C for seven days to identify plants with asynaptic chromosomes at metaphase I of meiosis. Two other Chinese Spring derived genotypes were used as controls for the KASP genotyping and in the temperature treatment experiments, the standard euploid (wild type) form, (AABBDD) and the nullisomic 5D-tetrasomic 5B (N5DT5B) genotype, which lacks chromosome 5D. Chinese Spring with homozygous *terminal* deletions of chromosome 5DL were obtained from TR Endo, Kyoto University, Japan. These were 5DL-1, 5DL-2, 5DL-5, 5DL-9 and 5DL-13 (Endo and Gill 1996).

### DNA extractions

Plants were initially grown in modular trays in a controlled environment room (CER) at 20°C (day) and 15°C (night) with a 16-hour photoperiod (lights on between 10:00 and 02:00) and 70 % humidity. Wheat seedlings were grown to the 2-3 leaf stage and approximately 5 cm of leaf material was harvested into 1.2 ml collection tubes containing 3 mm tungsten-carbide beads in a 96-well format on dry ice. DNA was extracted using the method of Chao and Somers https://maswheat.ucdavis.edu/protocols/general_protocols/DNA_extraction_003.htm (original reference in Pallotta et al. 2003) except that leaf material was shaken in a Geno/Grinder (Spex) at 1500 rpm for 2 min. Extracted DNA was diluted with dH_2_0 so that final DNA template concentrations were between 15-30 ng.

### Genotyping 5DL terminal deletion lines

Nine chromosome 5D specific microsatellite markers (Somers et al. 2004), from the GrainGenes database https://wheat.pw.usda.gov/GG3/ were used to map the breakpoints of the five Chinese Spring 5DL terminal deletion lines (Fig. 1). Following PCR amplification, products were separated by agarose gel electrophoresis.

**Fig. 1.**
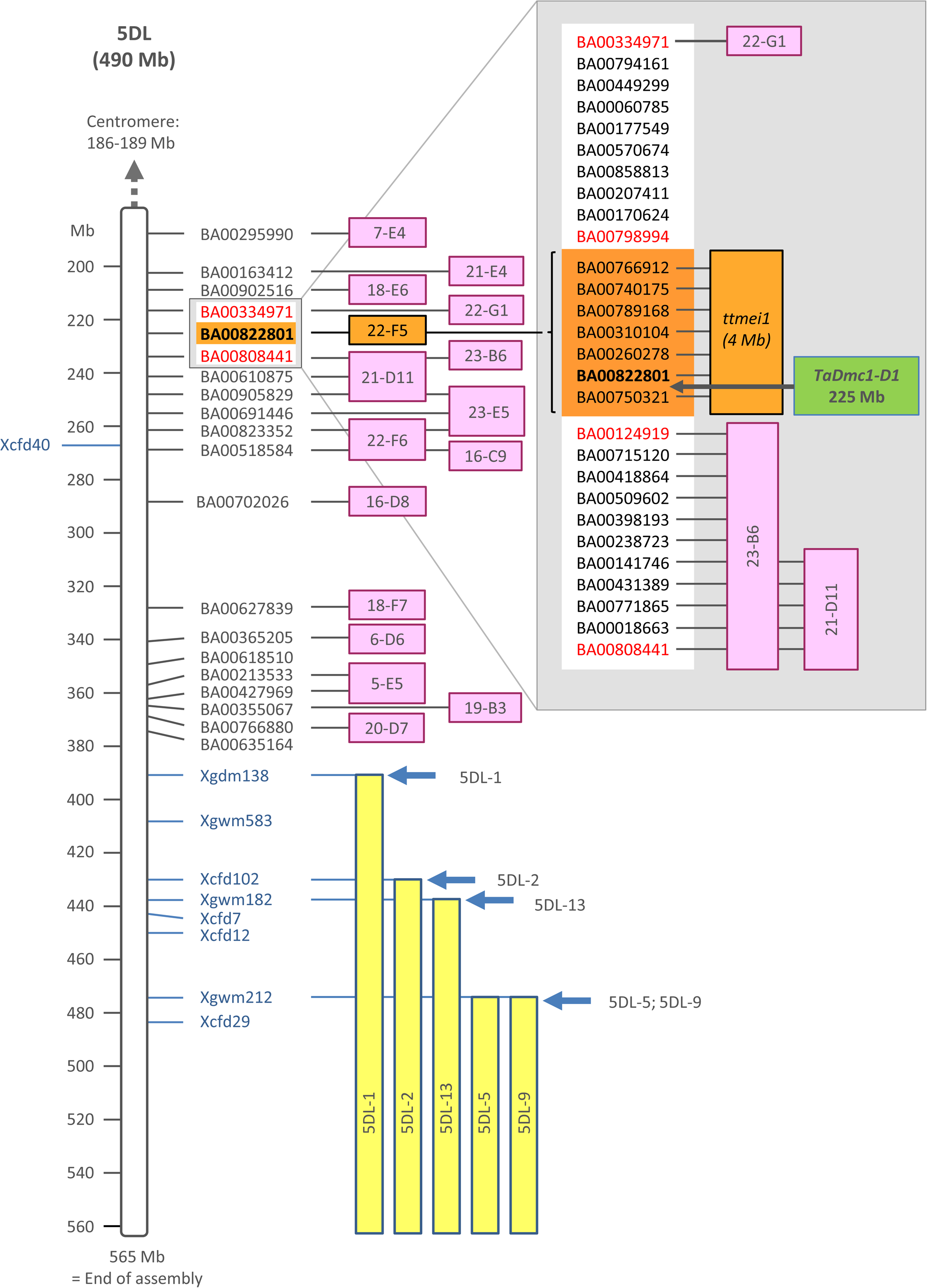
Map of wheat chromosome arm 5DL showing locations of deletions detected using 5DL specific KASP markers (black or red text with BA prefix) or microsatellite markers (blue); Markers are aligned with positions on the Chinese Spring IWGSC RefSeq v1.0 genome assembly (shown in Mb); Yellow boxes mark the extent of the 5DL terminal deletions in lines with normal pairing at 13°C; Breakpoints of terminal deletion lines are indicated by blue arrows; Pink boxes show positions of 5DL interstitial deletions in lines with normal pairing at 13°C; Orange boxes show the location of the 5DL deletion in the mutant line 22-F5 (*ttmei1)*, which exhibits asynapsis at 13°C; Small grey box shows the *ttmei1* deletion region and its flanking markers (in red) as detected in the initial KASP genotyping analysis with the only marker deleted in *ttmei1* (BA00822801) shown in bold text; Larger grey inset box shows fine mapping between flanking markers BA00334971 and BA00808441 (red text) using BA00822801 and 25 additional KASP markers. Markers shown in orange boxes are deleted in the *ttmei1* mutant. The position of the candidate gene *TaDmc1-D1* (green box) at 225 Mb is indicated with a black arrow

### KASP genotyping

KASP genotyping of M_2_ plants was performed using SNP (single nucleotide polymorphism) markers from the Wheat Breeders’ 820 k Axiom® array (Winfield et al. 2016) available at www.cerealsdb.uk.net. This dataset was aligned with the Chinese Spring reference sequence assembly, IWGSC RefSeq v1.0, (IWGSC 2018), using Python Data Matcher https://github.com/wingenl/python_data_matcher, which merges files with common features. The Biomart data mining tool from Ensembl Plants (Kinsella et al. 2011), available at http://plants.ensembl.org/biomart/martview/, was used as a source for the RefSeq v1.0 assembly coordinates.

Initially, KASP markers used for genotyping were selected at 15 Mb intervals along chromosome 5DL, between the centromere and the breakpoint of the most proximal Chinese Spring terminal deletion line. Later, more KASP markers were used to increase the density across regions of interest (Fig. 1). Selected KASP Primers were 5D chromosome-specific and had homeologous SNPs at the 3’ end. The allele-specific forward primers and common reverse primers were synthesised by Sigma-Aldrich. Allele-specific primers were synthesised with standard FAM or VIC compatible tails at their 5’ ends (FAM tail: 5’ GAAGGTGACCAAGTTCATGCT 3’; VIC tail: 5’ GAAGGTCGGAGTCAACGGATT 3’). Twenty primer sets were used in the first round of screening but following the identification of the *ttmei1* 5DL deletion mutant, 25 more KASP markers were selected between markers BA00334971 and BA00808441 to fine map the deletion (Fig. 1).

### KASP reaction and PCR conditions

The KASP reaction and its components were as recommended by LGC Genomics Ltd and described at https://www.biosearchtech.com/support/education/kasp-genotyping-reagents/how-does-kasp-work Assays were set up as 5 μl reactions in a 384-well format and included 2.5 μl genomic DNA template (15-30 ng of DNA), 2.5 μl of KASP 2x Master Mix (LGC Genomics), and 0.07 μl primer mix. Primer mix consisted of 12 μl of each tailed primer (100 μM), 30 μl common primer (100 μM) and 46 μl dH2O. PCR amplification was performed using an Eppendorf Mastercycler Pro 384 thermal cycler (Eppendorf, UK) using the following programme: Hotstart at 94°C for 15 min, followed by ten touchdown cycles (94°C for 20 s; touchdown from 65-57°C for 1 min, decreasing by 0.8°C per cycle) and then 30 cycles of amplification (94°C for 20 s; 57°C for 1 min). Fluorescent signals from PCR products were read in a PHERAstar microplate reader (BMG LABTECH Ltd.). If tight genotyping clusters were not obtained, an additional 5 cycles (94°C for 20 s; 57°C for 1 min) were performed. Genotyping data was analysed using KlusterCaller software (LGC Genomics).

### Low and high temperature treatments

Plants were initially grown in a CER at 20°C (day) and 15°C (night) with a 16-hour photoperiod (lights on between 10:00 and 02:00) and 70 % humidity until Zadoks growth stage 39 (Zadoks et al. 1974; Tottman 1987) when the flag leaf ligule is just visible. They were then transferred to plant growth cabinets under continuous light and exposed to low temperatures (13°C) for seven days (with 70 % humidity) or high temperatures (30°C) for 24 hours (75 % humidity). At 20°C, meiosis takes around 24 hours to complete (Bennett et al. 1971, 1973). However, high temperatures speed up meiosis and low temperatures slow it down (Bennett et al. 1972), so the different lengths of the high and low temperature treatments ensured that plants would be exposed to temperature treatments during the period from premeiotic interphase to early meiosis I, when they are most sensitive to temperature stress. It also ensured that the anthers to be sampled would have pollen mother cells (PMCs) containing chromosomes at metaphase I at the end of the treatment period so that synapsis and CO formation could be scored. At 30°C, 24 hours is long enough to ensure meiosis progresses to metaphase I but minimises the time available for adverse effects of high temperature on the plants. Longer periods of high temperature treatment would have made scoring PMCs more difficult as the chromosomes become sticky (Bayliss and Riley 1972a). To prevent dehydration during high temperature treatment, plant pots were kept in trays of water. For the high temperature treatments, plants were placed into the treatment cabinets at approximately the same time of day, between 10.00 and 11.45am.

### Preparation of PMCs for phenotyping

After temperature treatment, anthers were collected from spikes estimated to be undergoing meiosis (when the flag leaf had fully emerged, and the spike length was 4-6 cm long). Anthers were sampled from the first (oldest) 5 tillers only. Using an M80 stereo microscope (Leica Microsystems Ltd., Milton Keynes, UK), anthers were dissected from the two largest florets in each spikelet. Only anthers at metaphase I were scored, so, to determine the meiotic stage, one anther from each floret was stained with acetocarmine and squashed under a cover slip to extrude the PMCs, which were then examined using a DM2000 light microscope (Leica Microsystems). The three anthers within any floret are synchronised in meiotic development, so when PMCs with metaphase I chromosomes were identified, the two remaining anthers from the same floret were fixed in 3:1 (v/v) 100% ethanol:acetic acid, for cytological analysis. Anthers were incubated for at least 24 hours at 4°C before being transferred to 70% ethanol. Fixed anthers were washed with 0.1% sodium dodecyl sulphate for 3-5 minutes, and then hydrolysed with 1M hydrochloric acid for 10 min at 60°C. They were then Feulgen stained with Schiff’s reagent and squashed in 45% acetic acid. This allowed the chromosomes to be spread more widely to facilitate scoring of crossover. Images of metaphase I chromosomes were captured using a DM2000 microscope equipped with a DFC450 camera and controlled by LAS v4.4 system software (Leica Microsystems). For each cell, images were captured in up to 8 different focal planes to aid scoring.

### Cytological analysis of chromosome crossover

For each plant, 20-30 PMCs were blind scored from digital images. For each cell, the different meiotic chromosome configurations were counted. These were unpaired univalents (0 chiasmata), rod bivalents (1 chiasma), ring bivalents (2 chiasmata), trivalents (2–3 chiasmata), tetravalents (3 chiasmata) and pentavalents (4 chiasmata). Chiasma frequency per PMC was calculated separately for single and double chiasmata (see Fig. 2 for examples of the scored structures).

**Fig. 2.**
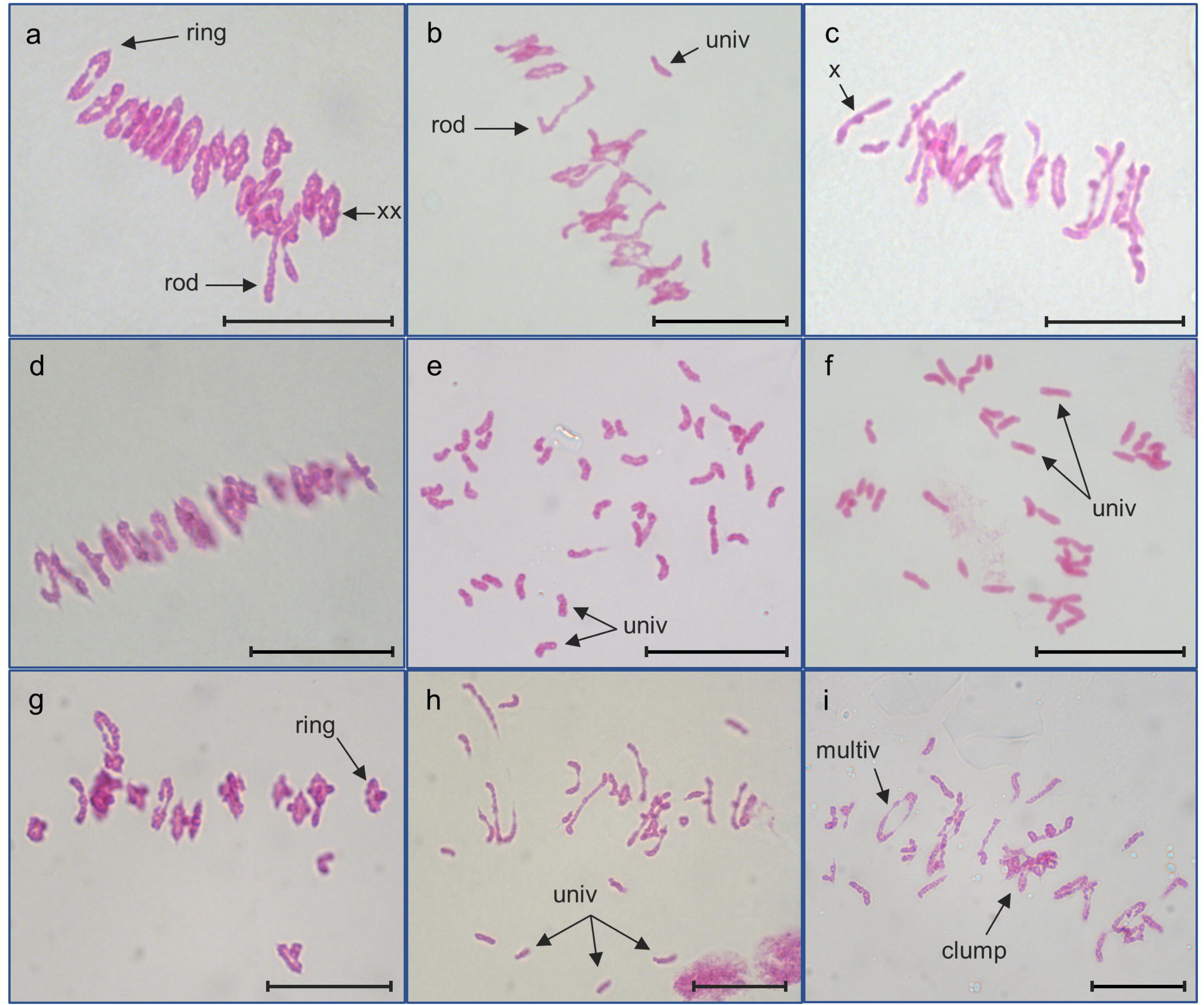
Feulgen stained metaphase I chromosomes from PMCs of wild-type Chinese Spring, *ttmei1* mutant and N5DT5B plants treated at different temperatures. a) wild type, b) *ttmei1* and c) N5DT5B at normal temperatures; d) wild type, e) *ttmei1* and f) N5DT5B after seven days at 13°C; g) wild type and h) and i) *ttmei1* after 24 hours at 30°C. Examples of univalent chromosomes (univ), rod bivalents (rod), ring bivalents (ring), multivalents (multiv), single chiasma (X), double chiasmata (XX) and sticky, clumping chromosomes (clump) are indicated with arrows; Note complete univalence in *ttmei1* and N5DT5B after treatment at 13°C. Scale bars, 10 μm

Statistical analyses were performed using STATISTIX 10.0 software (Analytical Software, Tallahassee, FL, USA). All treatments were analysed by the Kruskal–Wallis test (nonparametric one-way analysis of variance). Means were separated using the Dunn’s test with a probability level of 0.05. Statistical analysis was carried out between temperatures, and between genotypes. Bar charts were plotted using Microsoft Excel (2016).

### Immunolocalisation of meiotic proteins ASY1 and ZYP1

To determine when meiosis is disrupted in the low temperature experiments, wild-type and *ttmei1* mutant PMCs were immunolabelled with antibodies against the meiotic proteins ASY1 and ZYP1. PMCs from wild type Chinese Spring and the *ttmei1* mutant were embedded in acrylamide pads to preserve their 3D structure, and immunolocalisation of ASY1 and ZYP1 was performed as described previously (Martín et al. 2014, 2017). Anti-TaASY1 (Boden et al. 2009) raised in rabbit was used at a dilution of 1:250 and anti-HvZYP1 (Colas et al. 2016) raised in rat was used at a dilution of 1:200. Anti-rabbit Alexa Fluor® 488 and anti-rat Alexa Fluor® 568 (Invitrogen) were used as secondary antibodies.

### Image acquisition and analysis

Polyacrylamide-embedded PMCs were optically sectioned using a DM5500B microscope (Leica Microsystems) equipped with a Hamamatsu ORCA-FLASH4.0 camera and controlled by Leica LAS-X software v2.0. Z-stacks were deconvolved using Leica LAS-X software. Images were processed using Fiji, which is an implementation of ImageJ, a public domain program available from http://rsb.info.nih.gov/ij/ (Schneider et al. 2012).

### *Dmc1* sequence analysis

Multiple sequence alignment was carried out using ClustalX2 software (Thompson et al. 1997; Larkin et al. 2007). BLAST searches of the *Dmc1* sequences from other plant species against the Chinese Spring IWGSC RefSeq v1.0 sequence assembly (IWGSC 2018) revealed that there are three homeologs of the *Dmc1* gene in hexaploid wheat: TraesCS5A02G133000 (*TaDmc1-A1*) on chromosome 5A, TraesCS5B02G131900 (*TaDmc1-B1*) on 5B and TraesCS5D02G141200 (*TaDmc1-D1*) on 5D. The nucleotide sequences of the DNA, the coding sequences (CDs) and the promoter regions (including 1500 nucleotides downstream of the start codon) of the three *TaDmc1* homeologs were compared.

Multiple DMC1 amino-acid sequence alignment was also carried out between hexaploid wheat (Chinese Spring), barley (*Hordeum vulgare*) and rice (*Oryza sativa* Japonica). Gene IDs from Ensembl Plants are, for barley: HORVU5Hr1G040730.3 (*HvDMC1*) and for rice: Os12g0143800 (*OsDMC1A*) and Os11g0146800 (*OsDMC1B*).

## Results

### Phenotyping 5DL terminal deletion lines at low temperatures

To locate the gene(s) responsible for the *Ltp1* phenotype, we first needed to narrow down our region of interest on chromosome 5DL. To facilitate this, we exposed five Chinese Spring lines with homozygous terminal deletions of 5DL to 13°C for seven days. All five had normal chromosome pairing at metaphase I. The breakpoints of these deletion lines were mapped using nine 5D-specific microsatellite markers (Fig. 1). The mapping results were consistent with the order of the breakpoints on the C-banding maps produced by Endo and Gill (1996). Line 5DL-1 had the largest terminal deletion. The breakpoint of this deletion lies between the markers Xcfd40 and Xgdm138, the latter being the most proximal marker to be deleted in this line. We inferred from this that the gene responsible for the *Ltp1* phenotype must lie proximal to Xgdm138. BLASTing the Xgdm138 primer sequence against the Chinese Spring IWGSC RefSeq v1.0 sequence assembly revealed its location to be at approximately 391 Mb. The length of chromosome 5D is estimated to be around 565 Mb and the centromere lies at 185.6-188.7 Mb (IWGSC 2018), so we can infer that the length of 5DL is around 376 Mb. The length of the chromosome distal to Xgdm138 is 174 Mb, so almost half of the chromosome arm could be eliminated from our search to find the gene responsible for the *Ltp1* phenotype.

### Identification of a 5DL deletion line with abnormal crossover frequency

A total of 2,444 Chinese Spring M_2_ plants were genotyped, initially using 20 KASP markers located in the proximal half of 5DL. This identified 16 plants with deletions in 5DL (Fig. 1). These deletion lines were all derived from seed irradiated with the higher dose (300Gy) of gamma irradiation. The largest of these deletions (in the mutant line 21-D11) was estimated to be between 16-23 Mb.

All 16 deletion mutants were exposed to 13°C for seven days during premeiotic interphase to early meiosis I. Fifteen plants had normal synapsis at metaphase I, but one deletion mutant, 22-F5, exhibited almost complete crossover failure after low temperature treatment (Fig. 2e; Table 1; Fig. 3). This deletion mutant also exhibited reduced crossover frequency at 30°C (Table 1; Fig. 3). We renamed this deletion mutant ‘*ttmei1’* to reflect the fact that it has a deletion of an unknown gene ‘*TTmei1’* (*T*emperature *T*olerant *mei*osis *1*). As the *ttmei1* deletion mutant has an identical phenotype to that described for the *Ltp1* locus, it suggests that *Ltp1* is among the genes that have been deleted in this mutant.

**Fig. 3.**
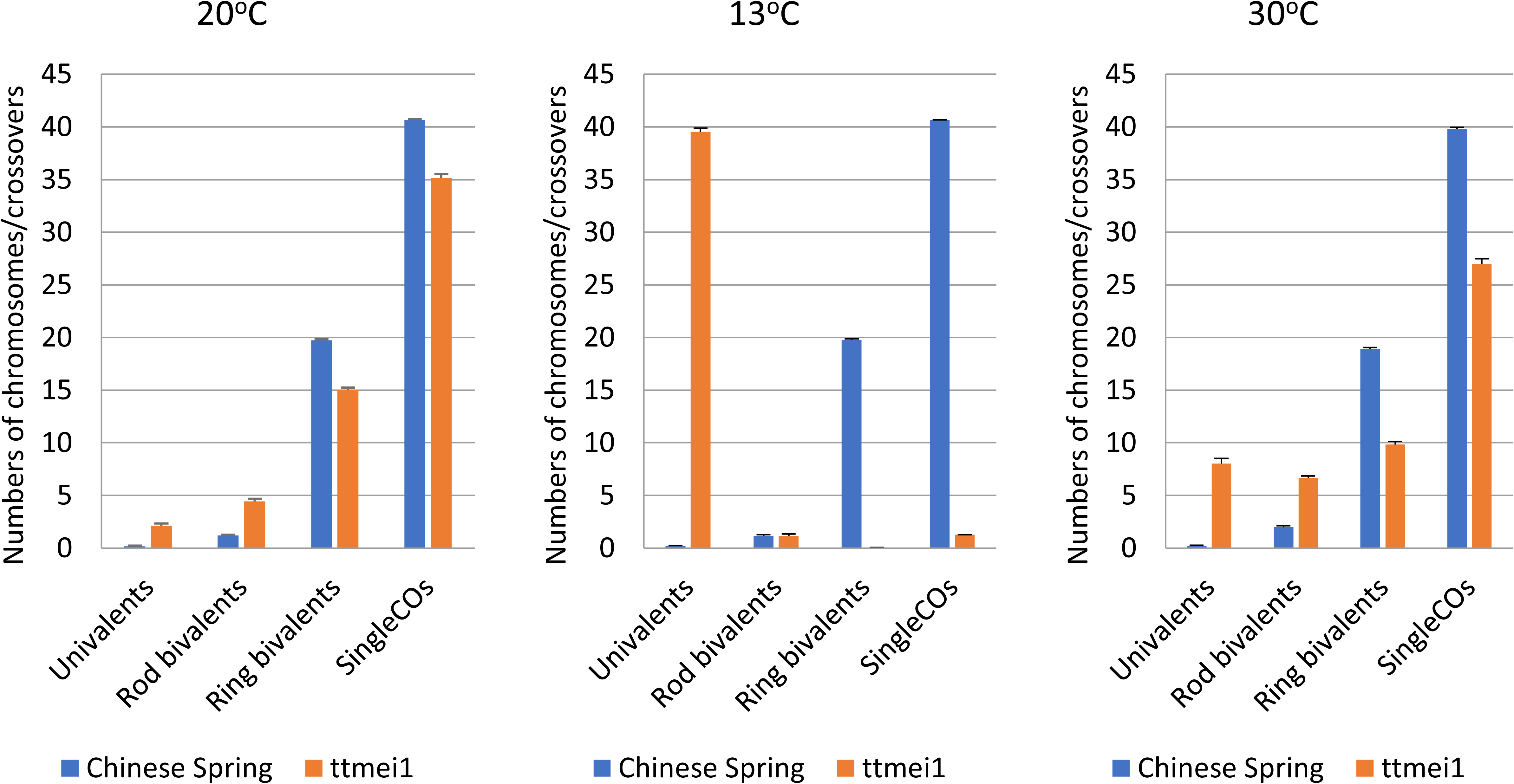
Bar charts showing genotypic effects on meiotic metaphase I chromosomes of Chinese Spring (CS) wild-type and *ttmei1* mutant plants after treatment at 20°C, 13°C and 30°C. The numbers of univalents, ring and rod bivalents and single crossovers are shown. Multivalents and double crossovers are not shown

**Table 1.**
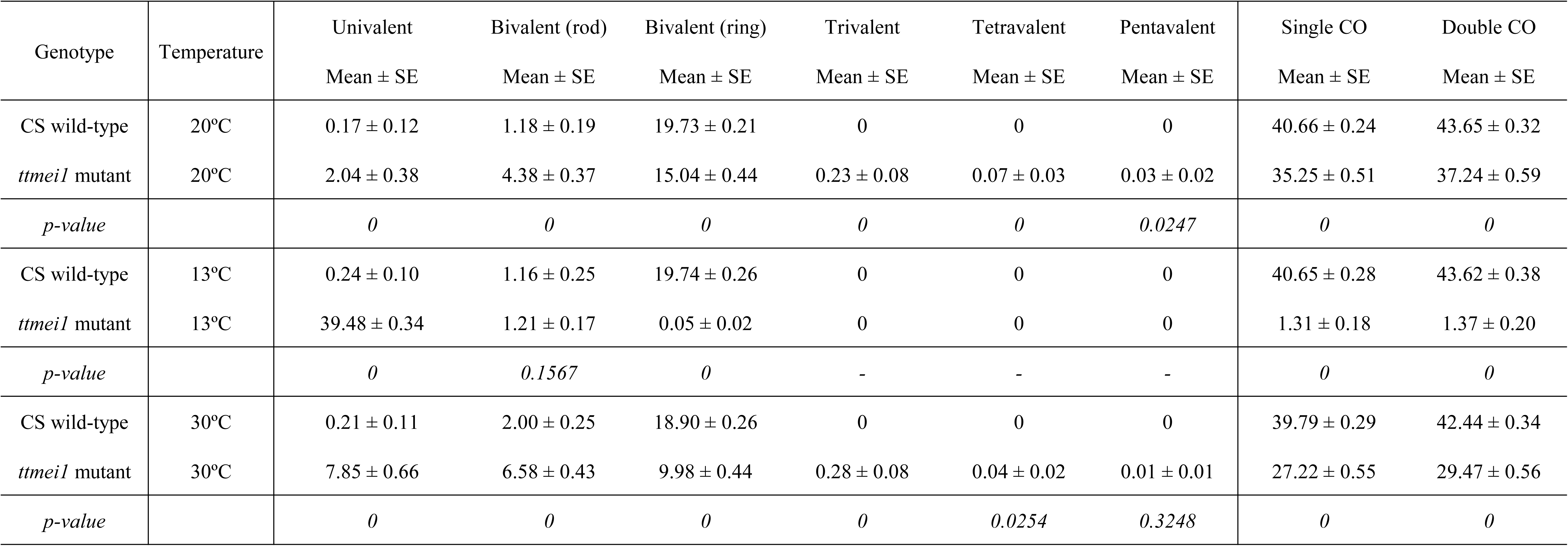
Genotypic effects on meiotic metaphase I chromosomes of Chinese Spring (CS) wild-type and *ttmei1* (22-F5) mutant plants after treatment at 20°C, 13°C and 30°C. The mean numbers of univalents, ring and rod bivalents and multivalents were scored along with chiasma frequency scored as single and double crossovers (CO). P-values < 0.05 indicate significant differences

### Crossover in *ttmei1* mutants at normal temperatures

The *ttmei1* mutant plant was self-fertilised and was able to produce M_3_ seed, although seed numbers appeared to be lower than that produced by wild-type plants. Further phenotypic analysis was carried out on Chinese Spring wild type and *ttmei1* M_3_ plants, which were grown under normal temperature conditions or were exposed to 13°C for seven days. Five plants were used for each treatment group. Statistical analysis identified differences in synapsis and crossover at different temperatures and between the two different genotypes.

At 20°C, in wild-type plants, all chromosomes lined up on the metaphase plate as normal, pairing as bivalents, mostly as rings but with the occasional rod bivalent (Fig. 2a). However, at the same temperature, the numbers of univalents, rod bivalents and multivalents were significantly higher in the *ttmei1* mutant when compared with the wild type, and the numbers of ring bivalents and both single and double crossovers were significantly lower (Table 1). Although the proportion of ring and rod bivalents was different in *ttmei1*, the mean number of bivalents (rings + rods) was around 19 as opposed to 21 in the wild type, and the chromosomes were still able to align on the metaphase plate with the exception of the occasional one or two univalent chromosomes (Fig. 2b). At 20°C, *ttmei1* chromosomes had similar conformations to those observed in N5DT5B plants (Fig. 2c).

### Crossover failure under low temperatures in *ttmei1* mutants

The 13°C treatment had no significant effect on wild-type metaphase I chromosomes (Table 2), which paired mainly as ring bivalents and lined up on the metaphase plate as for normal temperatures (Fig. 2a and d). However, in the *ttmei1* mutant, there were significantly higher numbers of univalent chromosomes at 13°C than at 20°C and significantly lower numbers of ring bivalents and crossovers (Table 2; Fig. 4). In *ttmei1* mutant plants at 13°C, the mean number of univalents per cell was almost 40, whereas in wild-type plants at the same temperature the mean number of univalents was less than one (Table 1; Fig. 3). In fact, in 60% of *ttmei1* PMCs *all* chromosomes were univalent. Univalents appeared more condensed than other chromosomes and were unable to align on the metaphase plate (Fig. 2e). The low chiasma frequency and high numbers of univalent chromosomes in *ttmei1* at low temperatures is identical in effect to that described when the whole of chromosome 5D is deleted in N5DT5B, as shown in Fig 2f.

**Fig. 4.**
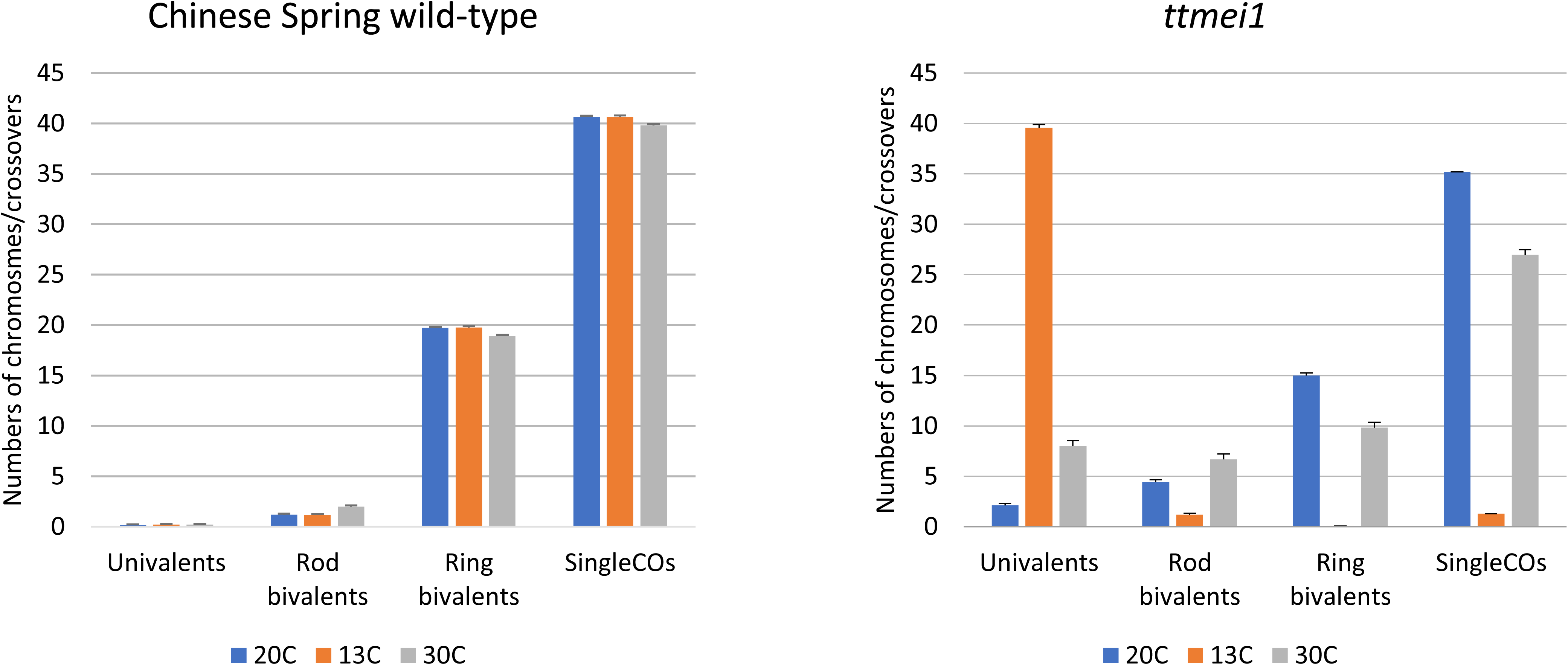
Bar charts showing the effects of three different temperature treatments (20°C, 13°C and 30°C) on meiotic metaphase I chromosomes of Chinese Spring (CS) wild-type and *ttmei1* mutant plants. The numbers of univalents, ring and rod bivalents and single crossovers are shown. Multivalents and double crossovers are not shown

**Table 2.** The effects of three different temperature treatments on meiotic metaphase I chromosomes of Chinese Spring (CS) wild-type and *ttmei1* mutant plants. The numbers of univalents, ring and rod bivalents and multivalents were scored along with chiasma frequency scored as single and double crossovers (CO). P-values < 0.05 indicate where there are significant differences in the scores between different temperature treatments. Superscript letters a, b and c indicate where the significant differences lie. For scores with the same letter, the difference between the means is not statistically significant. If the scores have different letters, they are significantly different

| Genotype | Temperature | Univalent | Bivalent (rod) | Bivalent (ring) | Trivalent | Tetravalent | Pentavalent | Single CO | Double CO |
| --- | --- | --- | --- | --- | --- | --- | --- | --- | --- |
|  |  | Mean ± SE | Mean ± SE | Mean ± SE | Mean ± SE | Mean ± SE | Mean ± SE | Mean ± SE | Mean ± SE |
| CS wild-type | 20°C | 0.17 ± 0.12 | 1.18 ± 0.19<sup>b</sup> | 19.73 ± 0.21<sup>a</sup> | 0 | 0 | 0 | 40.66 ± 0.24<sup>a</sup> | 43.65 ± 0.32<sup>a</sup> |
| CS wild-type | 13°C | 0.24 ± 0.10 | 1.16 ± 0.25<sup>b</sup> | 19.74 ± 0.26<sup>a</sup> | 0 | 0 | 0 | 40.65 ± 0.28<sup>a</sup> | 43.62 ± 0.38<sup>a</sup> |
| CS wild-type | 30°C | 0.21 ± 0.11 | 2.00 ± 0.25<sup>a</sup> | 18.90 ± 0.26<sup>b</sup> | 0 | 0 | 0 | 39.79 ± 0.29<sup>b</sup> | 42.44 ± 0.34<sup>b</sup> |
| *p-value* |  | 0.9964 | 0 | 0 | - | - | - | 0 | 0 |
| *ttmei1* mutant | 20°C | 2.04 ± 0.38<sup>c</sup> | 4.38 ± 0.37<sup>b</sup> | 15.04 ± 0.44<sup>a</sup> | 0.23 ± 0.08<sup>a</sup> | 0.07 ± 0.03<sup>a</sup> | 0.03 ± 0.02 | 35.25 ± 0.51<sup>a</sup> | 37.24 ± 0.59<sup>a</sup> |
| *ttmei1* mutant | 13°C | 39.48 ± 0.34<sup>a</sup> | 1.21 ± 0.17<sup>c</sup> | 0.05 ± 0.02<sup>c</sup> | 0<sup>b</sup> | 0<sup>c</sup> | 0 | 1.31 ± 0.18<sup>c</sup> | 1.37 ± 0.20<sup>c</sup> |
| *ttmei1* mutant | 30°C | 7.85 ± 0.66<sup>b</sup> | 6.58 ± 0.43<sup>a</sup> | 9.98 ± 0.44<sup>b</sup> | 0.28 ± 0.08<sup>a</sup> | 0.04 ± 0.02<sup>b</sup> | 0.01 ± 0.01 | 27.22 ± 0.55<sup>b</sup> | 29.47 ± 0.56<sup>b</sup> |
| *p-value* |  | 0 | 0 | 0 | 0 | 0.0247 | 0.1121 | 0 | 0 |

### Crossover failure under high temperatures in *ttmei1* mutants

Five wild-type and 11 *ttmei1* M_3_ plants were exposed to 30°C for 24 hours. A higher number of mutant plants were treated because of the wide variation in the data for this group. Chromosome stickiness and clumping was observed in many of the cells at this temperature (Fig. 2i), which made scoring more difficult than at normal or low temperatures. In wild-type plants, high temperature resulted in significantly higher numbers of rod bivalents and significantly lower numbers of ring bivalents and crossovers than had been observed at 20°C (Fig. 2g; Table 2; Fig. 4). However, these differences were much more pronounced in the *ttmei1* mutant. In *ttmei1* mutant plants there were significantly higher numbers of univalents, rod bivalents and tetravalents at 30°C than at normal temperatures, and significantly fewer ring bivalents and crossovers although the level of univalence was not nearly as pronounced as that at 13°C (Table 2; Fig. 4). There were also significant differences between the two genotypes at 30°C. For example, the mean number of univalents was 7.85 in *ttmei1* mutants but only 0.21 in wild-type plants. It was also observed that many *ttmei1* chromosomes did not align correctly on the metaphase plate (Fig. 2h and i).

### Immunolabelling of PMCs at low temperatures

To investigate when meiosis was being disrupted in the low temperature experiments, wild-type and *ttmei1* mutant PMCs were immunolabelled with antibodies against the meiotic proteins ASY1 and ZYP1 to follow the progression of synapsis (Fig. 5). ASY1 is part of the lateral elements of the SC, and first appears during premeiotic interphase, before synapsis begins (Armstrong et al. 2002; Boden 2009). ZYP1 is part of the central region of the SC, which assembles between the lateral elements (Higgins et al. 2005; Khoo et al 2012) and it is present only where chromosomes are synapsed. Immunolabelling therefore gives an indication of the level of synapsis achieved at pachytene when the SC is fully established, which enables tracking of the progression of synapsis. Clear differences were detected when comparing ASY1 (green) and ZYP1 (magenta) signals in the wild type and *ttmei1* mutant at low temperatures. In wild-type wheat (Fig. 5a), synapsis proceeds in the same way as it does at normal temperatures: It starts from the telomeres at one pole of the nucleus during early zygotene, and, by pachytene, all chromosomes have synapsed. A very small amount of ASY1 labelling is still visible at pachytene, indicating the small amount of chromatin that has not yet synapsed. In *ttmei1* mutants (Fig5b), at early zygotene, ASY1 localises to the chromosome axes in a pattern similar to that observed in wild-type wheat, and synapsis initiates normally at one pole of the nucleus. After early zygotene, chromatin staining with DAPI (4′ suggests that the *ttmei1* PMCs are at pachytene (Fig. 5b), but the amount of ZYP1 labelling is much lower than in pachytene in the wild-type, indicating that synapsis has been compromised and is not completed. None of the PMCs had a completed synapsis in the *ttmei1* mutant.

**Fig. 5.**
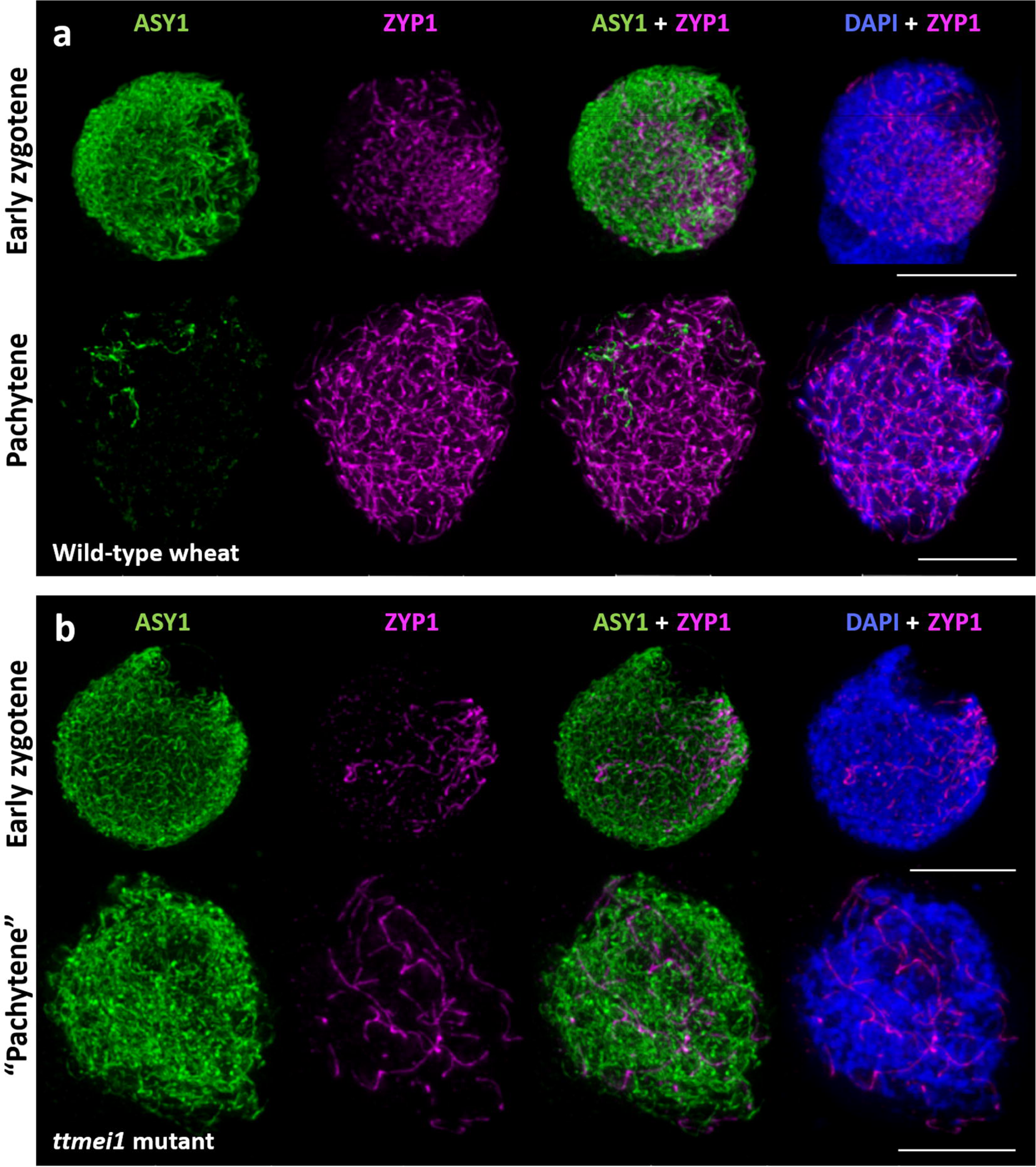
Immunolocalisation of meiotic proteins ASY1 (green) and ZYP1 (magenta) in PMCs from wheat Chinese Spring wild type (a) and *ttmei1* mutant (b), both under low temperature conditions

a. During early zygotene, synapsis starts from the telomeres at one pole of the nucleus in wild-type wheat. During pachytene, all chromosomes have synapsed. ASY1 labelling in green shows that a very small proportion of chromatin has not yet synapsed
b. In *ttmei1* mutants, synapsis initiates normally at one pole of the nucleus, however, synapsis is soon compromised and is not completed. Therefore, there is no normal pachytene in the *ttmei1* mutant. DAPI staining in blue. Scale bar, 10 μm

### Gene content in the *ttmei1* deletion region

The *ttmei1* deletion was initially mapped to a 16 Mb region of 5DL, between the flanking markers BA00334971 and BA00808441 (Fig. 1). In this initial analysis, only one KASP marker, BA00822801, was deleted in *ttmei1*. KASP genotyping of *ttmei1* using 25 more markers between BA00334971 and BA00808441, fine-mapped the deletion to a 4 Mb region between BA00798994 and BA00124919 (Fig. 1). Seven markers were deleted in this region.

The gene content within the 4 Mb deletion regions in *ttmei1* was revealed using data derived from the hexaploid wheat gene annotation v1.1 (IWGSC 2018) available from Ensembl Plants. Functional annotations of the genes were retrieved from the file “FunctionalAnnotation.rds” in https://opendata.earlham.ac.uk/wheat/under_license/toronto/Ramirez-Gonzalez_etal_2018-06025-Transcriptome-Landscape/data/TablesForExploration/FunctionalAnnotation.rds (Ramírez-González et al. 2018). Expression patterns of all the genes within the *ttmei1* deletion region were investigated based on 876 RNASeq samples from different tissue types available in the wheat expression browser (http://www.wheat-expression.com). A total of 41 genes (16 high confidence and 25 low confidence genes) were identified in the 4 Mb deletion region. Eighteen of the 41 genes were expressed during meiosis (Table 3). We considered a gene to be expressed when its expression level was > 0.5 transcripts per million (TPM) in at least one meiotic sample (Alabdullah et al. 2019). Twelve of these eighteen genes are high confidence genes. Detailed expression data (TPM) for all genes in the *ttmei1* deletion region is shown in Supplementary Table 1.

**Table 3.**
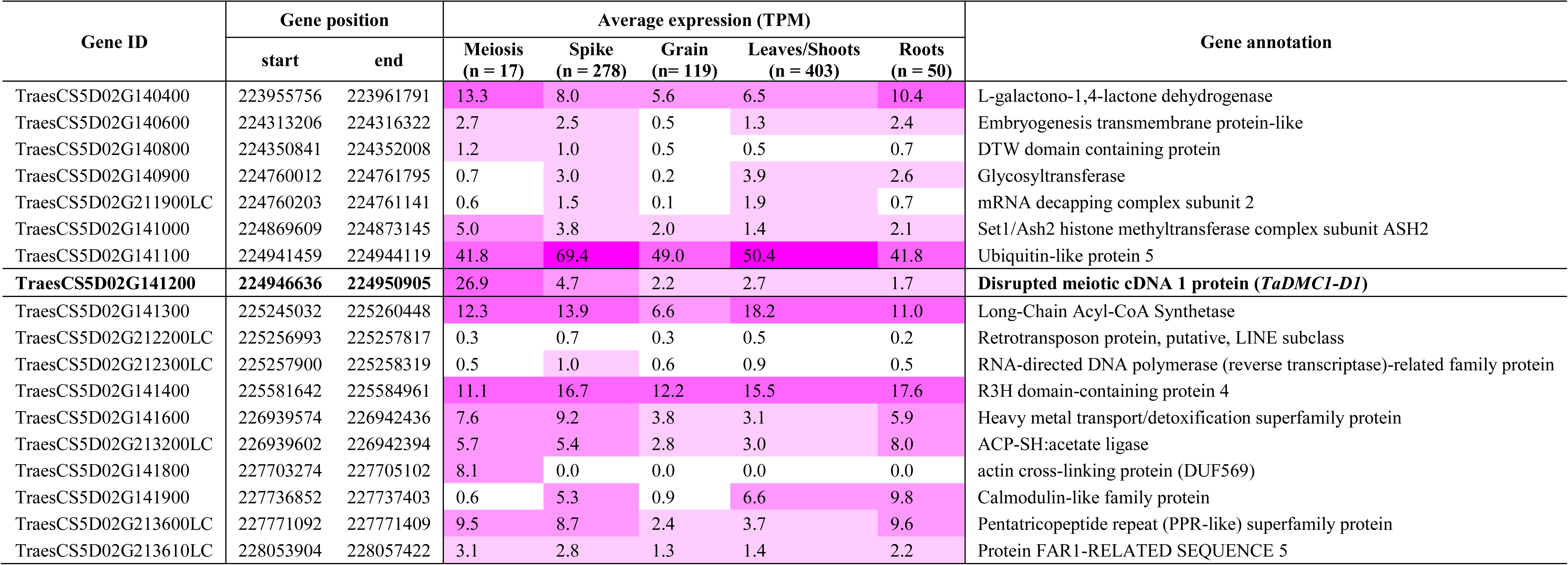
Genes expressed during meiosis within the 4 Mb deletion region of *ttmei1*. *TaDmc1-D1* is shown in bold. Both High Confidence and Low Confidence (LC) genes are included. A total of 867 RNA-Seq samples from different tissue types, including 17 meiotic anther samples, were used to determine the expression pattern of the genes. A gene was defined as being expressed during meiosis when its expression level > 0.5 TPM (transcripts per million) in at least one meiotic anther sample. Gene position is according to the Chinese Spring IWGSC RefSeq v1.1 gene annotation and was retrieved from Ensembl Plants

### *TaDmc1* is a candidate for the *Ltp1* phenotype

Of the 18 genes expressed during meiosis in the *ttmei1* deletion interval, the strongest candidate for the *TTmei1* phenotype is TraesCS5D02G141200, which is the D-genome homeolog of *Dmc1* (*Disrupted meiotic cDNA 1)*, which has a known meiotic phenotype, playing a central role in homologous recombination. It is a high confidence gene, and its expression level is around ten times higher in meiotically active tissues compared with non-meiotic tissues (averages 26.9 TPM and 2.8 TPM, respectively [Suppl. Fig. 1]). The other 17 genes in the region are much less likely than *TaDmc1* to be candidates for the *TTmei1* phenotype because none of them could be attributed to any known meiotic function, and, although they are expressed during meiosis, expression is proportionally higher in other tissues.

BLAST searches against the Chinese Spring IWGSC RefSeq v1.0 sequence assembly (IWGSC 2018) revealed that there are three homeologs of the *Dmc1* gene in hexaploid wheat: TraesCS5A02G133000 (*TaDmc1-A1*) on chromosome 5A, TraesCS5B02G131900 (*TaDmc1-B1*) on 5B and TraesCS5D02G141200 (*TaDmc1-D1*) on 5D. There are differences in gene expression levels between the three *Dmc1* homeologs in meiotic anthers: *TaDmc1-A1* has the lowest gene expression levels and *TaDmc1-D1* the highest (Suppl. Fig. 1).

### *Dmc1* sequence analysis

The nucleotide sequences of the DNA, the coding sequences (CDs) and the promoter regions of the three *TaDmc1* homeologs were compared. The CDs of the A, B and D homeologs are highly conserved (Suppl. Fig. 2) with sequence identity ranging between 98.1% and 98.7% (Suppl. Table 2). The DNA sequences are less similar (Suppl. Fig. 3), with sequence identity ranging from 82.1% (between A and B homeologs) to 89.0% (between A and D homeologs) (Suppl. Table 2). Comparing the promoter regions of each of the three *TaDmc1* homeologs showed a large insertion mutation (163 nt in size) at position 430 downstream of the start codon of *TaDmc1-B1* (Suppl. Fig. 4).

The exon-intron structure of the three *TaDmc1* homeologs (Fig. 6a) is conserved in terms of number of exons (14 exons each), however there is variation in intron length between the three homeologs due to large indel mutations mainly in introns 3, 6 and 7 (data not shown). *TaDmc1* homeologs have no splice variants. In all three *TaDmc1* homeologs, there is an open reading frame of 1035 bp (Suppl. Fig. 2), which encodes a predicted protein of 344 amino acids (Fig. 6b). This protein is highly conserved between the three homeologs. There are five single amino-acid substitutions between the wheat homeologs, but only one single amino-acid substitution (at position 114) in TaDMC1-D1 (threonine) when compared with TaDMC1-A1 and TaDMC1-B1 (alanine) (Fig. 6b).

**Fig. 6.**
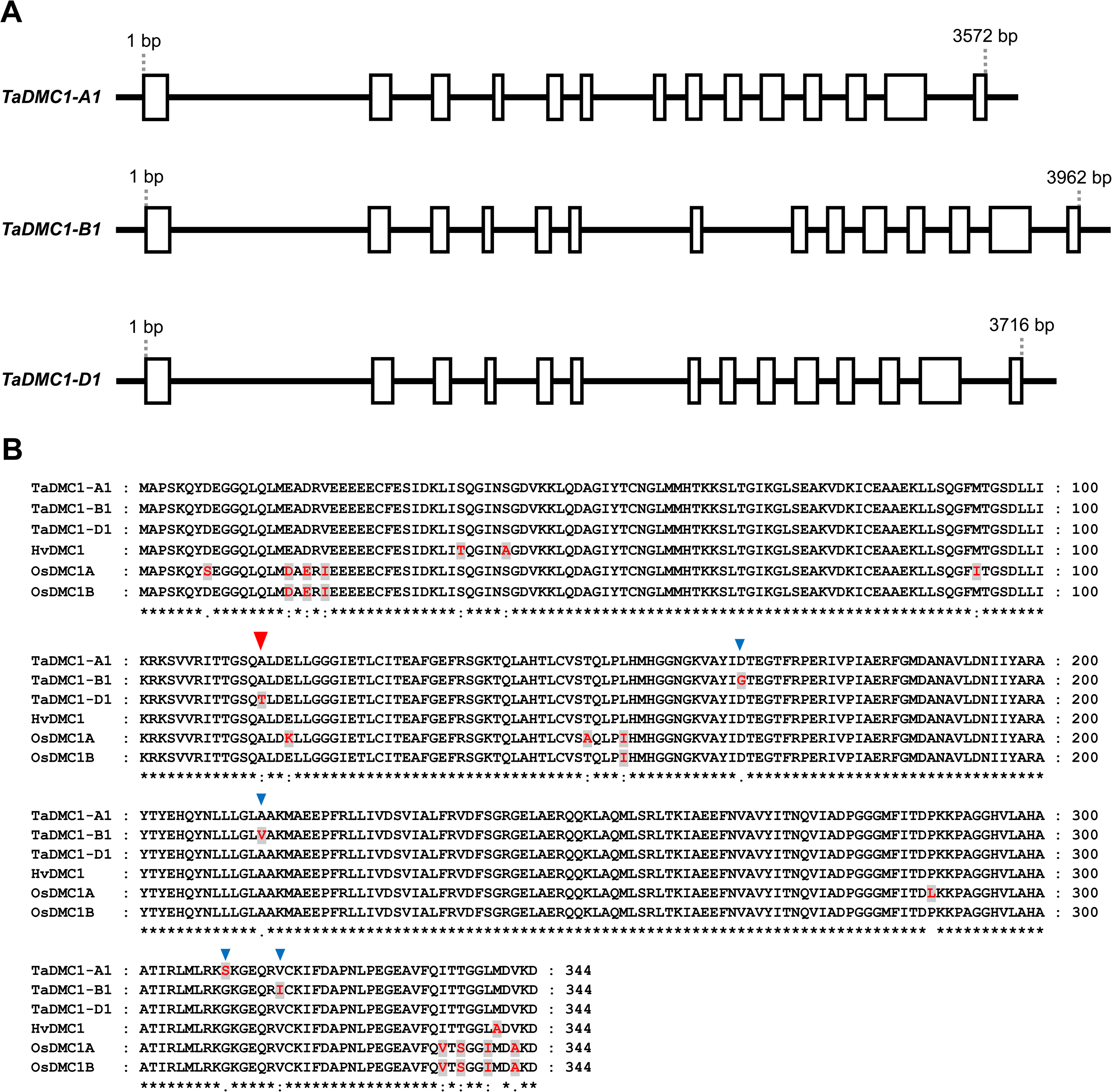
*TaDmc1* gene structure and amino-acid sequence alignment. (a) Exon-intron structure of the three *TaDmc1* homeologs (b) Multiple sequence alignment of amino-acids from DMC1 proteins of different cereal plants: hexaploid wheat (*Triticum aestivum* cv Chinese Spring; Ta), barley (*Hordeum vulgare*; Hv) and rice (*Oryza sativa* Japonica; Or). SNPs are shown in red text on a grey background. Symbols below each position in the sequence indicate the amount of conservation (asterisk “*”: identical residues; colon “:”: conserved substitution; period “.”: semi-conserved substitution; and space “ “: not conserved). A single amino-acid substitution in TaDMC1-D1 at position 114 (indicated by a red triangle) may confer low temperature tolerance in wheat. Blue triangles show positions of four other amino-acid substitutions in the TaDMC1-A1 and TaDMC1-B1 proteins

Multiple DMC1 amino-acid sequence alignment was also carried out between the three hexaploid wheat homeologs, barley (*Hordeum vulgare*) and rice (*Oryza sativa* Japonica). There is a high level of amino acid sequence conservation between the wheat, barley and rice proteins (Fig. 6b) with sequence identity ranging between 95.3 % and 99.4 % (Suppl. Table 2). This conservation of amino-acid sequence between the wheat, barley and rice DMC1 proteins has also been reported by Colas et al. (2019).

## Discussion

Chinese Spring wild-type plants and mutant plants with interstitial deletions within 5DL were exposed to seven days of low temperature (13°C) or 24 hours of high temperature (30°C) during the period from premeiotic interphase to early meiosis I. Subsequent microscopic examination of PMCs undergoing male meiosis enabled us to assess the effects of temperature on chromosome synapsis and crossover formation and to identify a candidate gene on 5DL (*TaDmc1-D1*) that is likely to be responsible for the *Ltp1* phenotype.

By exploiting a series of wheat 5DL terminal deletion lines, the *Ltp1* locus was initially delimited to the proximal half of chromosome 5DL. Using terminal deletion lines to narrow down our region of interest proved to be a good strategy as we were able to eliminate almost the entire distal half of the chromosome from our investigations. The deletion mapping was refined by screening a population of around 2,500 gamma irradiated plants using 5DL specific KASP markers. Of these, sixteen plants had 5DL deletions. The resources now available, including the Chinese Spring RefSeq V1.0 genomic sequence genome assembly, KASP primers from www.cerealsdb.uk.net and the Ensembl Plants database, made the screening and mapping processes much easier and quicker than when a similar approach was used to map the *Ph1* locus (Griffiths et al. 2006).

Exposure of the 5DL mutant plants to 13°C for seven days identified a deletion mutant, *ttmei1* that exhibits extremely high levels of chromosome univalence at low temperatures. This is identical to the *Ltp1* phenotype previously described in N5DT5B plants where the whole of chromosome 5D is missing. When grown under normal conditions, *ttmei1* plants are not sterile and are able to produce seed; however, seed numbers appeared to be lower than in the wild type. In *ttmei1* mutants, chromosome synapsis and crossover are only slightly disrupted at normal temperatures. However, the presence of high numbers of univalents and more rod bivalents than normal in *ttmei1* mutants under low temperatures, suggests a major problem with CO formation. Consistent with this, the *ttmei1* line exhibits significant abnormalities of synapsis at low temperatures.

In *ttmei1* mutants, aberrant chiasma formation was also observed after 24 hours at 30°C, but this was not as pronounced as in the low temperature phenotype. There is high variation in the scores for chiasma frequency between the *ttmei1* plants at 30°C. One explanation for this variation could be that plants were placed into the 30°C treatment cabinets at slightly different times of day (10.00-11.45am), so that high temperature treatment may have coincided with temperature sensitive stages of meiosis or premeiosis to a greater or lesser extent. It is also possible that *ttemi1* mutants carry a background deletion that has not yet been detected that contains a different gene affecting high temperature tolerance that is segregating with the *ttmei1* deletion. We are currently conducting further experiments to understand this variation at high temperature.

The *ttmei1*mutant carries a 4 Mb deletion of 5DL. Within this deletion interval there are 41 genes, 18 of which are expressed during meiosis. Amongst these genes, the strongest candidate for the low temperature pairing phenotype is the meiotic recombination gene *TaDmc1-D1*. None of the other genes are known to have any meiotic function. Moreover, *TaDmc1* is expressed most highly during early prophase I (Suppl. Table 1), which, in wheat, is when synapsis is initiated at the ‘telomere bouquet’ stage (Martín et al. 2017). *TaDmc1-D1* is therefore a strong candidate for temperature tolerance at low temperatures and, to a lesser extent, at high temperatures.

DMC1 plays a central role in homologous recombination, SC formation and cell-cycle progression. It is a meiosis-specific protein, structurally similar to the bacterial strand exchange protein RECA (Bishop et al. 1992). The control of DNA strand exchange during meiotic recombination is vital for sexual reproduction. Homologous recombination is initiated by the programmed formation of DNA double-strand breaks (DSBs) at leptotene. DMC1 and RAD51 (another RECA homolog) form filaments on the single-strand DNA overhangs at DSBs. These filaments facilitate homology search and catalyse strand invasion and strand exchange between homologous chromosomes (Neale and Keeney 2006). Repair of these interhomolog invasion events results in crossovers or non-crossovers (Reviewed in Lambing et al. 2017).

*Dmc1* homologs are found in a wide variety of organisms. In yeast, *dmc1* mutants have defects in reciprocal recombination, fail to form normal SCs, accumulate DSB recombination intermediates and the cells arrest in late meiotic prophase (Bishop et al. 1992). In mice, *dmc1* mutants have defective synapsis which leads to severe sterility also due to prophase arrest (Pittman et al. 1998; Yoshida et al. 1998; Bannister et al. 2007). In our wheat experiments, immunolocalisation of ASY1 and ZYP1 showed that at 13°C synapsis was compromised in the *ttmei1* mutant and was not completed, with meiosis appearing to arrest before pachytene in late prophase. This is similar to the phenotypes observed in yeast and mice at ambient temperatures.

In most diploid plant species, deletion of *Dmc1* also leads to sterility (Devisetty 2010). However, in *Arabidopsis thaliana*, *atdmc1* mutants have a disrupted synapsis with high levels of univalence, resulting in abnormal pollen grain formation and reduced fertility, but plants are not completely sterile because random chromosome segregation allows enough PMCs to reach maturity (Couteau et al. 1999). The Arabidopsis genome contains one copy of the *Dmc1* gene. In rice (*Oryza sativa*), there are two *Dmc1* homologs, *OsDMC1A* and *OsDMC1B*, that probably arose through chromosome duplication (Ding et al. 2001). A mutation in one or other of these homologs does not cause problems in meiosis, but *Osdmc1a Osdmc1b* double mutants exhibit serious CO defects, abnormal synapsis, high numbers of univalents at metaphase and are sterile (Wang et al. 2016). Barley (*Hordeum vulgare*) carries a single *Dmc1* homolog, *HvDMC1*, and mutations in this gene lead to abnormal synapsis, multiple univalents and chromosome mis-segregation (Colas et al. 2019; Szurman-Zubrzycka et al. 2019). Thus, it appears that disruption of the barley orthologue of *Dmc1* at normal temperatures leads to a phenotype similar to that of *ttmei1* at low temperatures.

There are three copies of *Dmc1* in bread wheat: *TaDmc1-A1* on 5A, *TaDmc1-B1* on 5B and *TaDmc1-D1 on* 5D. In the hexaploid wheat variety ‘Highbury’, there are differences in the levels of gene expression between the three *Dmc1* homeologs during premeiosis and early meiosis, which suggests that they might contribute to meiotic recombination to differing extents (Devisetty et al. 2010). This is reflected in our own study in Chinese Spring, where, in meiotic anthers, expression levels of *TaDmc1-D1* are higher than those of *TaDmc1-A1* and *TaDmc1-B1* (Suppl. Fig. 1).

In Chinese Spring, deletion of the region containing *TaDmc1-D1* results in asynapsis at 13°C and it seems highly likely that this gene promotes low temperature tolerance. In the absence of *TaDmc1-D1*, the *TaDmc1-A1* and *TaDmc1-B1* alleles are unable to stabilise synapsis at 13°C, at least when present as a single dose. There is only one amino-acid difference between the TaDMC1-D1 protein and the TaDMC1-A1 and TaDMC1-B1 proteins (Fig. 6b), so this substitution of alanine (a nonpolar amino acid) by threonine (a polar amino acid) at position 114 could account for the differences in temperature tolerance, although differences could also relate to expression effects such as arise from differences in the promoter regions (Suppl. Fig. 4).

Four other amino-acid differences between the 5A and 5B copies of TaDMC1 may be responsible for previously reported differences in temperature sensitivity when either chromosome 5A or 5B is missing. Previous studies suggest that chromosome 5A has a weak stabilising effect on chromosome pairing at low temperatures and, if present as a double dose, can compensate for a deficiency of chromosome 5D, whereas chromosome 5B cannot (Riley et al. 1966), so different dosage of the *TaDmc1* alleles may affect the stability of synapsis at low temperatures. It is important that breeders are aware that there could be *TaDmc1* alleles in their wheat germplasm that cannot stabilise wheat synapsis at extremes of temperature, and that the effects of these alleles may be masked by the temperature stabilising actions of TaDMC1-D1.

Interestingly, a previous study has shown that some other varieties and subspecies of wheat differ from Chinese Spring in that the gene that stabilises chromosome pairing at low temperatures is located on a chromosome other than 5D (Chapman and Miller 1980). In the wheat variety ‘Hope’ and the subspecies *spelta*, the *Ltp1* gene is located on chromosome 5A rather than 5D. It has also been postulated that there are additional *Ltp* genes on chromosomes 5AS (*Ltp2*) and 5BS (*Ltp3*) that also have a stabilising effect on homologous pairing and crossover at low temperatures (Queiroz et al. 1991), but high levels of univalence and low chiasma frequencies were only seen after 20 days of cold treatment and at a slightly lower temperature (10°C).

### Implications for plant breeding

In the future, breeders face the challenge of producing crops with increasing resilience to environmental stress whilst producing ever higher yields (Steinmeyer et al. 2013). Global climate models predict that temperatures will continue to increase over the 21st century and that high temperature extremes such as heat waves are likely to occur with a higher frequency and duration (Collins et al. 2013). It has been estimated that if global temperatures increase by just one degree (°C), wheat yields will decrease by 6% (Asseng et al. 2015). In the main European wheat-growing areas, the occurrence of adverse weather conditions might substantially increase by 2060 resulting in more frequent crop failure (Trnka et al. 2014). This could threaten global food security since Europe produces almost a third of the world’s wheat.

Moreover, the probability of multiple adverse events occurring within one season is projected to increase sharply by mid-century. Cold stress at meiosis could become more common in future years because climate change is expected to result in mild winters and warm springs, which are likely to speed up plant development prematurely, resulting in exposure of vulnerable plant tissues and organs to subsequent late season frosts. This occurred in 2007 when severe low temperatures following a period of above average temperatures caused widespread damage to agriculture in the United States (Gu et al. 2008). Models also predict an increase in frequency of heat stress at meiosis (Semenov et al. 2014). Our experiments have shown that even a relatively short period of high temperature at a critical stage of wheat development is sufficient for meiosis to be significantly affected. If current weather trends continue, it will become increasingly important to cultivate both heat tolerant varieties and cold tolerant varieties.

Heat tolerance is thought to be a complex trait in plants and is likely to be under the control of multiple genes (Barnabás et al. 2008), but we still know relatively little about the role of individual genes controlling temperature tolerance in wheat (Mullarkey and Jones 2000). Identification of *TaDmc1* as a candidate gene that can stabilise chromosome synapsis against extremes of temperature may allow wheat breeders to exploit this information and can provide markers that will allow them to identify and select hexaploid wheat genotypes that carry low (and high) temperature tolerance alleles at this locus.

## Acknowledgements

We are grateful to TR Endo for providing seed of the Chinese Spring 5DL terminal deletion lines; The International Atomic Energy Agency, Vienna for gamma irradiating the Chinese Spring seed; Philippa Borrill for the functional annotation data; Luzie Wingen for the Python Data matcher script for the alignment of datasets.

---

**Suppl. Fig. 1** Gene expression patterns of the three hexaploid wheat *TaDmc1* homeologs, TraesCS5A02G133000 (*TaDmc1-A1*) on chromosome 5A, TraesCS5B02G131900 (*TaDmc1-B1*) on 5B and TraesCS5D02G141200 (*TaDmc1-D1*) on 5D in different tissue types, based on the 876 RNASeq samples available in the wheat expression browser (http://www.wheat-expression.com). *TaDmc1* expression levels are higher in meiotically active tissues than non-meiotic tissues. In meiotic anthers, expression levels of *TaDmc1-D1* are higher than those of *TaDmc1-A1* and *TaDmc1-B1*

**Suppl. Fig. 2** Multiple alignment of the coding sequences (CDs) of the three hexaploid wheat *Dmc1* homeologs: TraesCS5A02G133000 (*TaDmc1-A1*) on chromosome 5A, TraesCS5B02G131900 (*TaDmc1-B1*) on 5B and TraesCS5D02G141200 (*TaDmc1-D1*) on 5D. SNPs are shown in red text on a grey background

**Suppl. Fig. 3** Multiple alignment of the DNA sequences of the three hexaploid wheat *Dmc1* homeologs: TraesCS5A02G133000 (*TaDmc1-A1*) on chromosome 5A, TraesCS5B02G131900 (*TaDmc1-B1*) on 5B and TraesCS5D02G141200 (*TaDmc1-D1*) on 5D

**Suppl. Fig. 4** Multiple sequence alignment of the promoter region of the three hexaploid wheat *Dmc1* homeologs: TraesCS5A02G133000 (*TaDmc1-A1*) on chromosome 5A, TraesCS5B02G131900 (*TaDmc1-B1*) on 5B and TraesCS5D02G141200 (*TaDmc1-D1*) on 5D. The promoter region includes the 1500 nucleotides downstream of the start codon of each of the *TaDmc1* homeologs. Note the large insertion mutation (163 nt in size) at position 430 downstream of the start codon of *TaDmc1-B1*

**Suppl. Table 1** Expression values (TPM) of all genes inside the *Dmc1* deletion region in 876 samples from different tissue types

**Suppl. Table 2** *Dmc1* sequence identity matrices. a) Percentages of amino acid sequence identity for DMC1 proteins in wheat (TaDMC1-A1, TaDMC1-B1 and TaDMC1-D1), barley (HvDMC1) and rice (OsDMC1A and OsDMC1B); b) Percentages of nucleotide sequence identity for the three *TaDmc1* homeologs TraesCS5A02G133000 (*TaDmc1-A1*) on chromosome 5A, TraesCS5B02G131900 (*TaDmc1-B1*) on 5B and TraesCS5D02G141200 (*TaDmc1-D1*) on 5D

## References

Alabdullah AK, Borrill P, Martin AC, Ramirez-Gonzalez RH, Hassani-Pak K, Uauy C, Shaw P, Moore G. (2019). A co-expression network in hexaploid wheat reveals mostly balanced expression and lack of significant gene loss of homeologous meiotic genes upon polyploidization. BioRxiv doi: https://doi.org/10.1101/695759

Armstrong SJ, Caryl AP, Jones GH, Franklin FCH (2002) Asy1, a protein required for meiotic chromosome synapsis, localizes to axis-associated chromatin in *Arabidopsis* and *Brassica*. J Cell Sci115: 3645–3655. https://doi.org/10.1242/jcs.00048

Asseng S, Ewert F, Martre P, Rotter RP, Lobell DB, Cammarano D, Kimball BA, Ottman MJ, Wall GW, White JW, Reynolds MP, Alderman PD, Prasad PVV, Aggarwal PK, Anothai J, Basso B, Biernath C, Challinor AJ, De Sanctis G, Doltra J, Fereres E, Garcia-Vila M, Gayler S, Hoogenboom G, Hunt LA, Izaurralde RC, Jabloun M, Jones CD, Kersebaum KC, Koehler AK, Muller C, Naresh Kumar S, Nendel C, O’Leary G, Olesen JE, Palosuo T, Priesack E, Eyshi Rezaei E, Ruane AC, Semenov MA, Shcherbak I, Stockle C, Stratonovitch P, Streck T, Supit I, Tao F, Thorburn PJ, Waha K, Wang E, Wallach D, Wolf J, Zhao Z, Zhu Y (2015) Rising temperatures reduce global wheat production. Nat Clim Change 5:143–147. https://doi.org/10.1038/nclimate2470

Bannister LA, Pezza RJ, Donaldson JR, de Rooij DG, Schimenti KJ, Camerini-Otero RD, Schimenti JC (2007) A Dominant, Recombination-Defective Allele of Dmc1 Causing Male-Specific Sterility. PLoS Biol 5: e105. https://doi.org/10.1371/journal.pbio.0050105

Barber HM, Carney J, Alghabari F, Gooding MJ (2015) Decimal growth stages for precision wheat production in changing environments? Ann Applied Biol 166:355–371

Barnabás B, Jäger K, Fehér A (2008) The effect of drought and heat stress on reproductive processes in cereals. Plant Cell Environ 31: 11–38. https://doi.org/10.1111/j.1365-3040.2007.01727.x

Bayliss MW, Riley R (1972a) An analysis of temperature-dependent asynapsis in *Triticum aestivum*. Genet Res 20: 193–200. https://doi.org/10.1017/S0016672300013707

Bayliss MW, Riley R (1972b) Evidence of premeiotic control of chromosome pairing in *Triticum aestivum*. Genet Res 20: 201–212

Bennett MD, Chapman V, Riley R (1971) The duration of meiosis in pollen mother cells of wheat, rye and *Triticale*. Proc R Soc Lond B 178: 259–275

Bennett MD, Rao MK, Smith JB, Bayliss MW (1973) Cell development in the anther, the ovule, and the young seed of *Triticum aestivum* L. var. Chinese Spring. Phil Trans R Soc B 266: 39–81

Bennett MD, Smith JB, Kemble R (1972) The effect of temperature on meiosis and pollen development in wheat and rye. Can J Genet Cytol 14: 615–624

Bilgir C, Dombecki CR, Chen PF, Villeneuve AM, Nabeshima K (2013) Assembly of the synaptonemal complex is a highly temperature-sensitive process that is supported by PGL-1 during *Caenorhabditis elegans* meiosis. G3 (Bethesda) 3: 585–595. https://doi.org/10.1534/g3.112.005165

Bishop DK, Park D, Xu L, Kleckner N (1992) *DMC1*: a meiosis-specific yeast homolog of E. coli *recA* required for recombination, synaptonemal complex formation, and cell cycle progression. Cell 69: 439–456. https://doi.org/10.1016/0092-8674(92)90446-J

Boden SA, Langridge P, Spangenberg G, Able JA (2009) TaASY1 promotes homologous chromosome interactions and is affected by deletion of *Ph1*. Plant J 57: 487–497

Bolser D, Staines DM, Pritchard E, Kersey P (2016) Ensembl Plants: Integrating Tools for Visualizing, Mining, and Analyzing Plant Genomics Data. In: Edwards D. (eds) Plant Bioinformatics. Methods in Molecular Biology, vol 1374. Humana Press, New York, NY

Bomblies K, Higgins JD, Yant L (2015) Meiosis evolves: adaptation to external and internal environments. New Phytol, 208: 306–323. https://doi.org/10.1111/nph.13499

Chapman V, Miller TE (1981) The location of a gene affecting meiotic chromosome pairing at low temperature in *Triticum aestivum*. Z. Pflanzenzüchtg 86: 50–55

Colas I, Barakate A, Macaulay M, Schreiber M, Stephens J, Vivera S, Halpin C, Waugh, Ramsay L (2019) desynaptic5 carries a spontaneous semi-dominant mutation affecting Disrupted Meiotic cDNA 1 in barley. J Exp Bot 70: 2683–2698. https://doi.org/10.1093/jxb/erz080

Colas I, Macaulay M, Higgins JD, Phillips D, Barakate A, Posch M, Armstrong SJ, Franklin FCH, Halpin C, Waugh R, Ramsay L (2016) A spontaneous mutation in MutL-homolog 3 (HvMLH3) affects synapsis and crossover resolution in the barley desynaptic mutant des10. New Phytol 212: 693–707. https://doi.org/10.1111/nph.14061

Collins, M., R. Knutti, J. Arblaster, J.-L. Dufresne, T. Fichefet, P. Friedlingstein, X. Gao, W.J. Gutowski, T. Johns, G. Krinner, M. Shongwe, C. Tebaldi, A.J. Weaver and M. Wehner, 2013: Long-term Climate Change: Projections, Commitments and Irreversibility. In: Climate Change 2013: The Physical Science Basis. Contribution of Working Group I to the Fifth Assessment Report of the Intergovernmental Panel on Climate Change [Stocker, T.F., D. Qin, G.-K. Plattner, M. Tignor, S.K. Allen, J. Boschung, A. Nauels, Y. Xia, V. Bex and P.M. Midgley (eds.)]. Cambridge University Press, Cambridge, United Kingdom and New York, NY, USA

Couteau F, Belzile F, Horlow C, Grandjean O, Vezon D, Doutriaux M-P (1999) Random chromosome segregation without meiotic arrest in both male and female meiocytes of a *dmc1* mutant of Arabidopsis. Plant Cell 11: 1623–1634. https://doi.org/10.1105/tpc.11.9.1623

De Storme N, Geelen D (2014) The impact of environmental stress on male reproductive development in plants: Biological processes and molecular mechanisms. Plant Cell Environ 37: 1–18

Devisetty UK (2010) Molecular investigation of RAD51 and DMC1 homoeologous genes of hexaploid wheat (*Triticum aestivum* L.). PhD thesis, University of Nottingham. http://eprints.nottingham.ac.uk/13340/1/523039.pdf

Devisetty UK, Mayes K, Mayes S. (2010). The *RAD51* and *DMC1* homoeologous genes of bread wheat: cloning, molecular characterization and expression analysis. BMC Res Notes 3: 245

Ding Z, Wang T, Chong K, Bai S (2001) Isolation and characterization of *OsDMC1*, the rice homologue of the yeast *DMC1* gene essential for meiosis. Sex Plant Reprod 13: 285. https://doi.org/10.1007/s004970100065

Dowrick GJ (1957) The influence of temperature on meiosis. Hered 11: 37–49. https://doi.org/10.1038/hdy.1957.4

Draeger T, Moore G (2017) Short periods of high temperature during meiosis prevent normal meiotic progression and reduce grain number in hexaploid wheat (*Triticum aestivum* L.). Theor Appl Genet 130: 1785–1800. https://doi.org/10.1007/s00122-017-2925-1

Elliott CG (1955) The effect of temperature on chiasma frequency. Hered 9: 385–398

Endo TR, Gill BS (1996) The Deletion Stocks of Common Wheat. J Hered 87: 295–307. https://doi.org/10.1093/oxfordjournals.jhered.a023003

Fischer RA (1985) Number of kernels in wheat crops and the influence of solar radiation and temperature. J Agr Sci 105:447–461. https://doi.org/10.1017/S0021859600056495

Fischer RA, Maurer R (1976) Crop temperature modification and yield potential in a dwarf spring wheat. Crop Sci 16: 855–859

Griffiths S, Sharp R, Foote TN, Bertin I, Wanous M, Reader S, Colas I, Moore G (2006) Molecular characterization of *Ph1* as a major chromosome pairing locus in polyploid wheat. Nature 439: 749–752. https://doi.org/10.1038/nature04434

Gu L, Hanson PJ, Post WM, Kaiser DP, Yang B, Nemani R, Pallardy SG, Meyers T (2008) The 2007 Eastern US Spring Freeze: Increased Cold Damage in a Warming World? BioScience 58: 253–262. https://doi.org/10.1641/B580311

Hayter AM (1969) Cytogenetics and cytochemistry of wheat species. PhD thesis. Cambridge University, Cambridge, England

Hayter AM, Riley R (1967) Duplicate genetic activities affecting meiotic chromosome pairing at low temperatures in *Triticum*. Nature 216: 1028–1029. https://doi.org/10.1038/2161028a0

Higgins JD, Perry RM, Barakate A, Ramsay L, Waugh R, Halpin C, Armstrong SJ, Franklin FCH (2012) Spatiotemporal asymmetry of the meiotic program underlies the predominantly distal distribution of meiotic crossovers in barley. Plant Cell 24: 4096–4109. https://doi.org/10.1105/tpc.112.102483

Higgins JD, Sanchez-Moran E, Armstrong SJ, Jones GH, Franklin FCH (2005). The Arabidopsis synaptonemal complex protein ZYP1 is required for chromosome synapsis and normal fidelity of crossing over. Genes Dev 19: 2488–2500

International Wheat Genome Sequencing Consortium (2018) Shifting the limits in wheat research and breeding using a fully annotated reference genome. Science 361: eaar7191 https://doi.org/10.1126/science.aar7191

Ji H, Xiao L, Xia Y, Song H, Liu B, Cao W, Zhu Y, Liu L (2017) Effects of jointing and booting low temperature stresses on grain yield and yield components in wheat. Agric For Meteorol 243: 33–42

Khoo KHP, Able AJ & Able JA (2012) The isolation and characterisation of the wheat molecular ZIPper I homologue, *Ta* ZYP1. BMC Res Notes 5: 106. https://doi.org/10.1186/1756-0500-5-106

Kinsella RJ, Kähäri A, Haider S, Zamora J, Proctor G, Spudich G, Almeida-King J, Staines D, Derwent P, Kerhornou A, Kersey P, Flicek P (2011). Ensembl BioMarts: a hub for data retrieval across taxonomic space. Database: 2011, bar030. https://doi.org/10.1093/database/bar030

Lambing C, Franklin FCH, Wang C-J R (2017) Understanding and Manipulating Meiotic Recombination in Plants. Plant Physiol 173: 1530–1542. https://doi.org/10.1104/pp.16.01530

Larkin MA, Blackshields G, Brown NP, Chenna R, McGettigan PA, McWilliam H, Valentin F, Wallace IM, Wilm A, Lopez R, Thompson JD, Gibson TJ, Higgins DG (2007) Clustal W and Clustal X version 2.0. Bioinformatics 23: 2947–2948

Law CN (1999) Sterility in winter wheat: review of occurrence in different varieties and possible causes. HGCA Res Review 44: 1–108

Martín AC, Shaw P, Phillips D, Reader S, Moore G (2014) Licensing MLH1 sites for crossover during meiosis. Nat Commun 5: 1–5

Martín AC, Rey M-D, Shaw P, Moore G (2017) Dual effect of the wheat *Ph1* locus on chromosome synapsis and crossover. Chromosoma 126: 669–680. https://doi.org/10.1007/s00412-017-0630-0

Mullarkey M, Jones P (2000) Isolation and analysis of thermotolerant mutants of wheat. J Exp Bot 51: 139–146. https://doi.org/10.1093/jexbot/51.342.139

Neale MJ, Keeney S (2006) Clarifying the mechanics of DNA strand exchange in meiotic recombination. Nature 442: 153–158

Page SL, Hawley RS (2004) The genetics and molecular biology of the synaptonemal complex. Ann Rev Cell Dev Biol 20: 525–558

Pallotta MA, Warner P, Fox RL, Kuchel H, Jefferies SJ, Langridge (2003) Marker assisted wheat breeding in the southern region of Australia. Proc 10th Int Wheat Genet Symp 2: 789–791

Pittman DL, Cobb J, Schimenti KJ, Wilson LA, Cooper DM, Brignull E, Handel MA, Schimenti JC (1998) Meiotic prophase arrest with failure of chromosome synapsis in mice deficient for *Dmc1*, a germline-specific RecA homolog. Mol Cell 1: 697–705

Porter JR, Gawith M (1999) Temperatures and the growth and development of wheat: a review. Eur J Agron10:23–36. https://doi.org/10.1016/S1161-0301(98)00047-1

Qin D, Wu H, Peng H, Yao Y, Ni Z, Li Z, Zhou C, Sun Q (2008) Heat stress-responsive transcriptome analysis in heat susceptible and tolerant wheat (*Triticum aestivum* L.) by using Wheat Genome Array. BMC Genomics 9: 432. https://doi.org/10.1186/1471-2164-9-432

Queiroz A, Mello-Sampayo T, Viegas WS (1991) Identification of low temperature stabilizing genes, controlling chromosome synapsis or recombination, in short arms of chromosomes from the homoeologous group 5 of *Triticum aestivum*. Hereditas 115: 37–41. https://doi.org/10.1111/j.1601-5223.1991.tb00344.x

Ramírez-González RH, Borrill P, Lang D, Harrington SA, Brinton J, Venturini L, Davey M, Jacobs J, van Ex F, Pasha A, Khedikar Y, Robinson SJ, Cory AT, Florio T, Concia L, Juery C, Schoonbeek H, Steuernagel B, Xiang D, Ridout CJ, Chalhoub B, Mayer KFX, Benhamed M, Latrasse D, Bendahmane A, International Wheat Genome Sequencing Consortium, Wulff BBH, Appels R, Tiwari V, Datla R, Choulet F, Pozniak CJ, Provart NJ, Sharpe AG, Paux E, Spannagl M, Bräutigam A, Uauy C (2018) The transcriptional landscape of polyploid wheat. Science 361: 662. https://doi.org/10.1126/science.aar6089

Riley R (1966) Genotype-environmental interaction affecting chiasma frequency in *Triticum aestivum*. In: Darlington CD, Lewis KR (eds) Chromosomes Today vol 1, Oliver & Boyd, Edinburgh. pp57–64

Riley R, Chapman V, Young RM, Bellfield AM (1966) Control of meiotic chromosome pairing by the chromosomes of homoeologous group 5 of *Triticum aestivum*. Nature 212: 1475–1477

Roberts MA, Reader SM, Dalgliesh C, Miller TE, Foote TN, Fish LJ, Snape JW, Moore G (1999) Induction and Characterization of *Ph1* Wheat Mutants. Genetics 153: 1909–1918

Saini HS, Aspinall D (1982) Abnormal sporogenesis in wheat (*Triticum aestivum* L.) induced by short periods of high temperature. Ann Bot 49: 835–846

Schneider CA, Rasband WS, Eliceiri KW (2012) NIH Image to ImageJ: 25 years of image analysis. Nat Methods 9:671–675

Semenov MA, Stratonovitch P, Alghabari F, Gooding MJ (2014) Adapting wheat in Europe for climate change. J Cereal Sci 59: 245–256. https://doi.org/10.1016/j.jcs.2014.01.006

Somers DJ, Isaac P, Edwards K (2004) A high-density wheat microsatellite consensus map for bread wheat (Triticum aestivum L.) Theor Appl Genet 109: 1105–1114. https://doi.org/10.1007/s00122-004-1740-7

Steinmeyer FT, Lukac M, Reynolds MP, Jones HE (2013) Quantifying the relationship between temperature regulation in the ear and floret development stage in wheat *(Triticum aestivum L.)* under heat and drought stress. Funct Plant Biol 40: 700–707. https://doi.org/10.1071/FP12362

Szurman-Zubrzycka M, Brygida B, Stolarek-Januszkiewicz M, Kwaś D (2019) The dmc1 mutant allows an insight into the DNA double-strand break repair during meiosis in barley (*Hordeum vulgare* L.). Front Plant Sci 10: 761 https://doi.org/10.3389/fpls.2019.00761

Thakur P, Kumar S, Malik JA, Berger JD, Nayyar H (2010) Cold stress effects on reproductive development in grain crops: An overview. Environ Exp Bot 67: 429–443. https://doi.org/10.1016/j.envexpbot.2009.09.004

Thompson JD, Gibson TJ, Plewniak F, Jeanmougin F, Higgins DG (1997) The ClustalX windows interface: flexible strategies for multiple sequence alignment aided by quality analysis tools. Nucleic Acids Res 25:4876–4882

Tottman DR (1987) The decimal code for the growth stages of cereals, with illustrations. Ann Appl Biol 110: 441–454. https://doi.org/10.1111/j.1744-7348.1987.tb03275.x

Trnka M, Rötter RP, Ruiz-Ramos M, Kersebaum KC, Olesen JE, Žalud Z, Semenov MA (2014) Adverse weather conditions for European wheat production will become more frequent with climate change. Nat Clim Change 4: 637–643. https://doi.org/10.1038/NCLIMATE2242

Wang H, Hu Q, Tang D, Liu X, Du G, Shen Y, Li Y, Cheng Z (2016) OsDMC1 is not required for homologous pairing in rice meiosis. Plant Physiol 171: 230–241. https://doi.org/10.1104/pp.16.00167

Wardlaw IF, Dawson IA, Munibi P, Fewster R (1989) The tolerance of wheat to high temperatures during reproductive growth. I. Survey procedures and general response patterns. Aust J Agric Res 40: 1–13. https://doi.org/10.1071/AR9890001

Winfield MO, Allen AM, Burridge AJ, Barker GLA, Benbow HR, Wilkinson PA, Coghill J, Waterfall C, Davassi A, Scopes G, Pirani A, Webster T, Brew F, Bloor C, King J, West C, Griffiths S, King I, Bentley AR, Edwards KJ (2016) High density SNP genotyping array for hexaploid wheat and its secondary and tertiary gene pool. Plant Biotech J 14:1195–1206 10.1111/pbi.12485

Yoshida K, Kondoh G, MatsudaY, Habu T, Nishimune Y, Morita T (1998) The mouse *RecA*-like gene *Dmc1* is required for homologous chromosome synapsis during meiosis. Molecular Cell 1: 707–718

Zadoks JC, Chang TT, Konzak CF (1974) A decimal code for the growth stages of cereals. Weed Research 14: 415–421

