## Supplementary figures and images for "*Dmc1* is a candidate for temperature tolerance during wheat meiosis"

### Supplemental Fig. 1

#### Slide 1
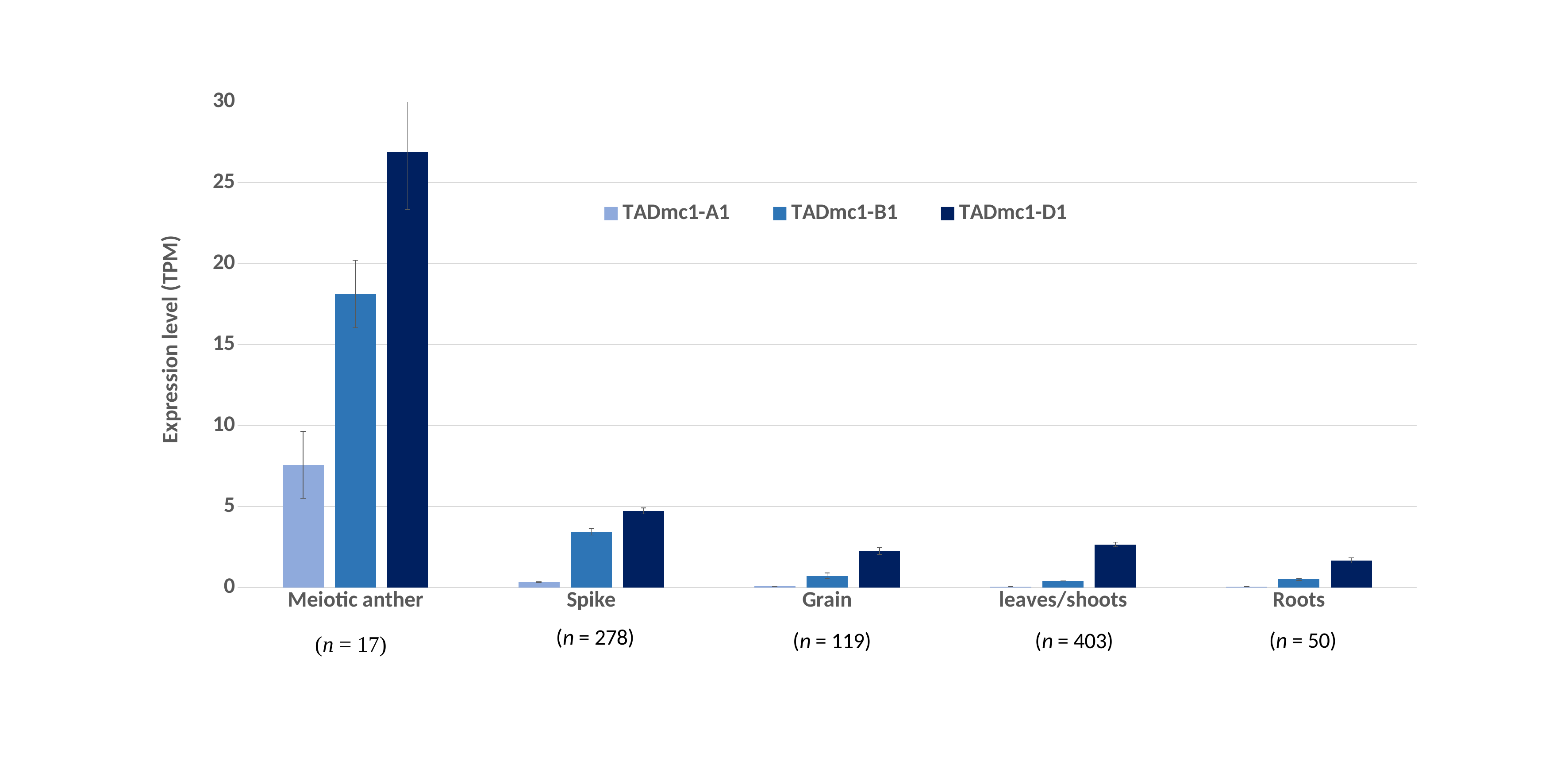

##### Chart
| Category | TADmc1-A1 | TADmc1-B1 | TADmc1-D1 |
|---|---|---|---|
| Meiotic anther | 7.573768588235294 | 18.12801 | 26.89158235294118 |
| Spike | 0.3441505607913666 | 3.432835413669065 | 4.73039065827338 |
| Grain | 0.06864552436974791 | 0.7136077403361344 | 2.2488975462184873 |
| leaves/shoots | 0.04916433066997518 | 0.41174925086848646 | 2.6522696714640177 |
| Roots | 0.049425460000000004 | 0.4994268979999999 | 1.6713086000000006 |
