## Supplemental Fig. 2 for "*Dmc1* is a candidate for temperature tolerance during wheat meiosis"

### Slide 1
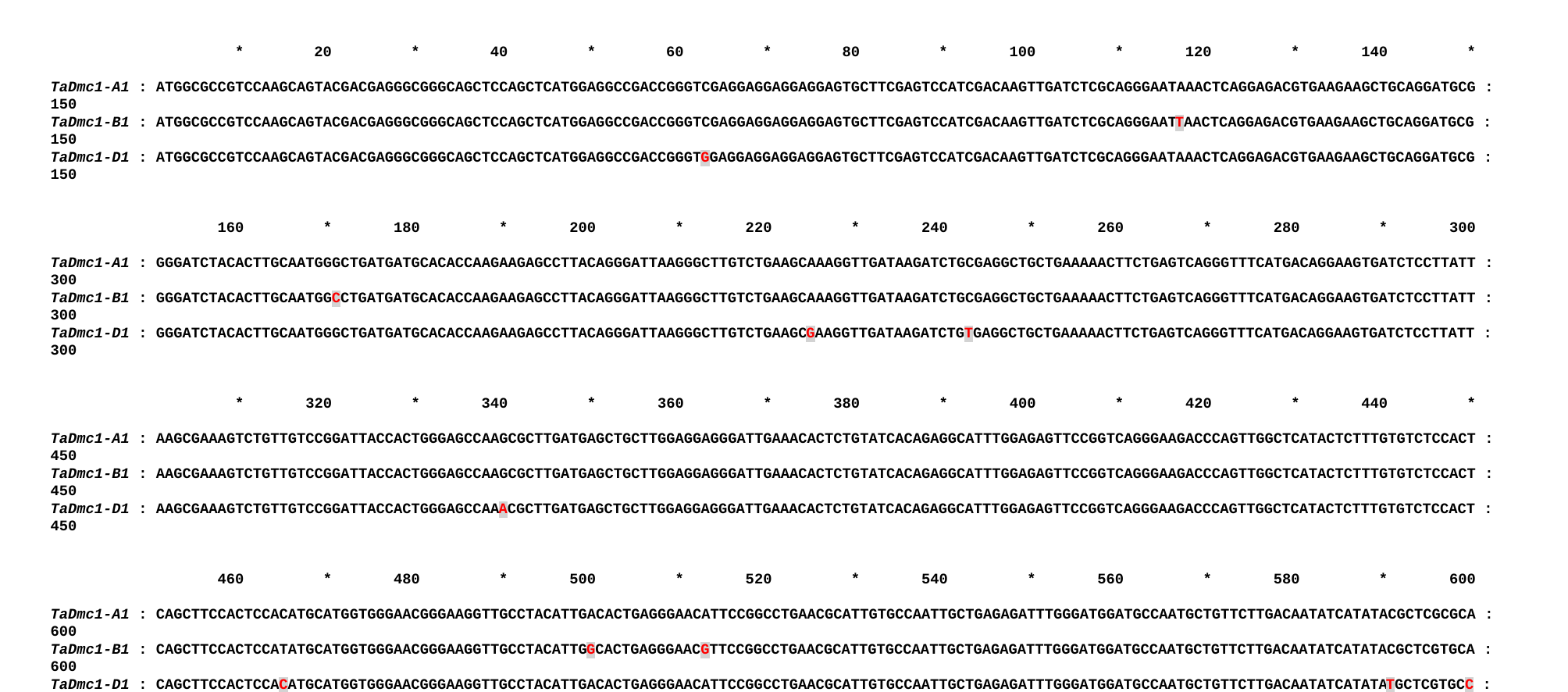

* 20 * 40 * 60 * 80 * 100 * 120 * 140 *
TaDmc1-A1 : ATGGCGCCGTCCAAGCAGTACGACGAGGGCGGGCAGCTCCAGCTCATGGAGGCCGACCGGGTCGAGGAGGAGGAGGAGTGCTTCGAGTCCATCGACAAGTTGATCTCGCAGGGAATAAACTCAGGAGACGTGAAGAAGCTGCAGGATGCG : 150
TaDmc1-B1 : ATGGCGCCGTCCAAGCAGTACGACGAGGGCGGGCAGCTCCAGCTCATGGAGGCCGACCGGGTCGAGGAGGAGGAGGAGTGCTTCGAGTCCATCGACAAGTTGATCTCGCAGGGAATTAACTCAGGAGACGTGAAGAAGCTGCAGGATGCG : 150
TaDmc1-D1 : ATGGCGCCGTCCAAGCAGTACGACGAGGGCGGGCAGCTCCAGCTCATGGAGGCCGACCGGGTGGAGGAGGAGGAGGAGTGCTTCGAGTCCATCGACAAGTTGATCTCGCAGGGAATAAACTCAGGAGACGTGAAGAAGCTGCAGGATGCG : 150
 160 * 180 * 200 * 220 * 240 * 260 * 280 * 300
TaDmc1-A1 : GGGATCTACACTTGCAATGGGCTGATGATGCACACCAAGAAGAGCCTTACAGGGATTAAGGGCTTGTCTGAAGCAAAGGTTGATAAGATCTGCGAGGCTGCTGAAAAACTTCTGAGTCAGGGTTTCATGACAGGAAGTGATCTCCTTATT : 300
TaDmc1-B1 : GGGATCTACACTTGCAATGGCCTGATGATGCACACCAAGAAGAGCCTTACAGGGATTAAGGGCTTGTCTGAAGCAAAGGTTGATAAGATCTGCGAGGCTGCTGAAAAACTTCTGAGTCAGGGTTTCATGACAGGAAGTGATCTCCTTATT : 300
TaDmc1-D1 : GGGATCTACACTTGCAATGGGCTGATGATGCACACCAAGAAGAGCCTTACAGGGATTAAGGGCTTGTCTGAAGCGAAGGTTGATAAGATCTGTGAGGCTGCTGAAAAACTTCTGAGTCAGGGTTTCATGACAGGAAGTGATCTCCTTATT : 300
 * 320 * 340 * 360 * 380 * 400 * 420 * 440 *
TaDmc1-A1 : AAGCGAAAGTCTGTTGTCCGGATTACCACTGGGAGCCAAGCGCTTGATGAGCTGCTTGGAGGAGGGATTGAAACACTCTGTATCACAGAGGCATTTGGAGAGTTCCGGTCAGGGAAGACCCAGTTGGCTCATACTCTTTGTGTCTCCACT : 450
TaDmc1-B1 : AAGCGAAAGTCTGTTGTCCGGATTACCACTGGGAGCCAAGCGCTTGATGAGCTGCTTGGAGGAGGGATTGAAACACTCTGTATCACAGAGGCATTTGGAGAGTTCCGGTCAGGGAAGACCCAGTTGGCTCATACTCTTTGTGTCTCCACT : 450
TaDmc1-D1 : AAGCGAAAGTCTGTTGTCCGGATTACCACTGGGAGCCAAACGCTTGATGAGCTGCTTGGAGGAGGGATTGAAACACTCTGTATCACAGAGGCATTTGGAGAGTTCCGGTCAGGGAAGACCCAGTTGGCTCATACTCTTTGTGTCTCCACT : 450
 460 * 480 * 500 * 520 * 540 * 560 * 580 * 600
TaDmc1-A1 : CAGCTTCCACTCCACATGCATGGTGGGAACGGGAAGGTTGCCTACATTGACACTGAGGGAACATTCCGGCCTGAACGCATTGTGCCAATTGCTGAGAGATTTGGGATGGATGCCAATGCTGTTCTTGACAATATCATATACGCTCGCGCA : 600
TaDmc1-B1 : CAGCTTCCACTCCATATGCATGGTGGGAACGGGAAGGTTGCCTACATTGGCACTGAGGGAACGTTCCGGCCTGAACGCATTGTGCCAATTGCTGAGAGATTTGGGATGGATGCCAATGCTGTTCTTGACAATATCATATACGCTCGTGCA : 600
TaDmc1-D1 : CAGCTTCCACTCCACATGCATGGTGGGAACGGGAAGGTTGCCTACATTGACACTGAGGGAACATTCCGGCCTGAACGCATTGTGCCAATTGCTGAGAGATTTGGGATGGATGCCAATGCTGTTCTTGACAATATCATATATGCTCGTGCC : 600
 * 620 * 640 * 660 * 680 * 700 * 720 * 740 *
TaDmc1-A1 : TACACCTATGAGCACCAGTACAACTTACTCCTGGGCCTTGCTGCCAAGATGGCCGAAGAGCCTTTCAGGCTTCTGATCGTGGATTCTGTGATTGCGCTATTCCGTGTTGATTTCAGTGGTAGGGGTGAACTTGCAGAGCGTCAGCAAAAA : 750
TaDmc1-B1 : TACACCTATGAGCACCAGTACAACTTACTCCTGGGCCTTGTTGCCAAGATGGCTGAAGAGCCTTTCAGGCTTCTGATCGTGGATTCTGTGATTGCGCTATTCCGTGTTGATTTCAGTGGCAGGGGTGAACTTGCAGAGCGTCAGCAAAAA : 750
TaDmc1-D1 : TACACCTATGAGCACCAGTACAACTTACTCCTGGGCCTTGCTGCCAAGATGGCTGAAGAGCCTTTCAGGCTTCTGATCGTGGATTCTGTGATTGCGCTGTTCCGTGTTGATTTCAGTGGTAGGGGTGAACTTGCAGAGCGTCAGCAAAAG : 750
 760 * 780 * 800 * 820 * 840 * 860 * 880 * 900
TaDmc1-A1 : CTGGCACAAATGCTGTCCCGCCTTACAAAGATTGCTGAGGAGTTCAATGTTGCAGTGTACATCACCAACCAAGTGATTGCGGACCCAGGTGGTGGTATGTTCATCACTGACCCCAAAAAGCCGGCAGGAGGCCACGTGCTGGCGCATGCA : 900
TaDmc1-B1 : CTGGCACAAATGCTTTCCCGCCTTACAAAGATTGCTGAGGAGTTCAATGTTGCAGTGTACATCACCAACCAAGTGATTGCGGACCCAGGTGGTGGTATGTTCATCACTGACCCCAAAAAGCCGGCGGGAGGCCACGTGCTGGCGCATGCA : 900
TaDmc1-D1 : CTGGCACAAATGCTGTCCCGCCTTACAAAGATTGCTGAGGAGTTCAATGTTGCAGTGTACATCACCAACCAAGTGATTGCGGACCCAGGTGGTGGTATGTTCATCACTGACCCCAAAAAGCCGGCGGGAGGCCACGTGCTGGCGCATGCA : 900
 * 920 * 940 * 960 * 980 * 1000 * 1020 *
TaDmc1-A1 : GCCACCATCCGGTTGATGCTGAGGAAAAGCAAAGGCGAGCAGCGTGTCTGCAAGATCTTTGACGCCCCTAACCTTCCCGAGGGAGAAGCTGTTTTCCAGATTACAACAGGCGGATTGATGGATGTGAAAGACTGA : 1035
TaDmc1-B1 : GCCACCATCCGGTTGATGCTGAGGAAAGGCAAAGGCGAGCAGCGTATCTGCAAGATCTTTGACGCCCCTAACCTTCCCGAGGGAGAAGCTGTTTTCCAGATTACAACAGGTGGATTGATGGATGTGAAAGACTGA : 1035
TaDmc1-D1 : GCCACCATCCGGTTGATGCTGAGGAAAGGCAAAGGCGAGCAGCGTGTCTGCAAGATCTTTGACGCCCCTAACCTTCCCGAGGGAGAAGCTGTTTTCCAGATCACAACAGGCGGATTGATGGATGTGAAAGACTGA : 1035
