## Supplemental Fig. 3 for "*Dmc1* is a candidate for temperature tolerance during wheat meiosis"

```
                     *        20         *        40         *        60         *        80         *       100         *       120         *       140         *
TaDmc1-A1 : ATGGCGCCGTCCAAGCAGTACGACGAGGGCGGGCAGCTCCAGCTCATGGAGGCCGACCGGGTCGAGGAGGAGGAGGAGTGCTTCGAGTCCATCGACAAGTGTACGTTCGCCGCCTCCTACCCCTCTCCTCTCGAAACCCGCCCGCTGCCC :  150
TaDmc1-B1 : ATGGCGCCGTCCAAGCAGTACGACGAGGGCGGGCAGCTCCAGCTCATGGAGGCCGACCGGGTCGAGGAGGAGGAGGAGTGCTTCGAGTCCATCGACAAGTGTATGTTCGCCGCCTCCAACTCCTCTCCTCTCGAAACCCTCCCGCCGCCC :  150
TaDmc1-D1 : ATGGCGCCGTCCAAGCAGTACGACGAGGGCGGGCAGCTCCAGCTCATGGAGGCCGACCGGGTGGAGGAGGAGGAGGAGTGCTTCGAGTCCATCGACAAGTGTACGTTCGCCGCCTCCAACCCCTCTCCTCTCCAAACCCTCCCGCCGCCC :  150

                   160         *       180         *       200         *       220         *       240         *       260         *       280         *       300
TaDmc1-A1 : CGTCTCCGGTGCTGCATTTACTTGCTTGTTCGTGTGCCTGCGTCGCGCGTGTGTCGGCATGTGGGGGTTAGGCCTGCTCACCGTTGCGTTCCGGGTGCTTCCGCCTCTAAGTTCGCGCGTTTCGGTCGCAATTTCGTGCTGTTTGGAGAT :  300
TaDmc1-B1 : CGTCTCCGGTGCTGCATTTACTTGCTTGTCCGTGTGCCTGCGTCGCGCGCGTGTCGGCGCGTGGGGGTTAGGCCTGCTCACCGTTGCGTTCCGGGTGCTTCTGCCTCTAAGTTCGCGCGTTTCGGTCACAATTTCGTGCTGTTTGGAGAT :  300
TaDmc1-D1 : CGTCTCCGGTGCTGCATTTACTTGCTTGTTCGTGTGCCTGCGTCGCGCGCGTGTCGGCGCGTGGGGGTTAGGCCTGCTCACCGTTGCGTCCCGGGTGCTTCCGCCTCTAAGTTCGCGCGTTTCGGTCGCAATTTCGTGCTGTTTGGAGAT :  300

                     *       320         *       340         *       360         *       380         *       400         *       420         *       440         *
TaDmc1-A1 : GGGTTTGGTGCGGATTTCGCTTAGCCTCCACATTTGGTTGGTTTTTGGTGCGTGCAGCGCGTGGTCGCGTGTGCTCGTGTTGCGTTTGTTTTGATTTTCTCCCCTATTTCGTGCAGTCTGGGGATTAACTAGCGCCCTT-GGTAGGCTAA :  449
TaDmc1-B1 : GGATTTGGTGCGGATTTCGCTTAGCCTCCACATTTGGTTGGTTTTTGGTGCGTGCAGCGCGTGGTCG--TGTGCTCGTGTTGCGTTGGTTTTGATTTTCTCCCCTATTTCGTGCAATCTGGGGATTAACTAGCGCCCTTTGGTAGCCTAA :  448
TaDmc1-D1 : GGATTTGGTGCGGATTTCGCTTAGCCTCCACATTTGGTTGGTTTTTGGTGCGTGCAGCGCGTGGTCG--TGTGCTCCTGTTGCGTTGGTTTTGATTTTCTCCCCTATGTCGTGCAATCTGGGGATTAACTAGCGCCCTTTGGTAGCCTAA :  448

                   460         *       480         *       500         *       520         *       540         *       560         *       580         *       600
TaDmc1-A1 : TTTCGTGCAGCTTTGCTGATTCCCCCCACTTGATTTCTCTTTACCGGTTAGTTTGTGCTAATCGCTGTGTGGTGCCGTTTTTTCTCCACGGATTAGTCTGAACTTAGCCCCAATTTTGAGCTGCCATGCGTTTAATCCGCCGTAGCGATC :  599
TaDmc1-B1 : TTTCGTGCGGCTTTTCTAATTCCCCCCACTTGATTTCTCTTTACCGGCTAGTTTGTGCTAATCGCTGTGTGGTGCCGTTTTTTCTCCACGGATTAGTCTGAACTTAGCCCCAATTTTGAGCTGCCATGCCTTTAATCCGCCGTATCGATC :  598
TaDmc1-D1 : TTTCGTGCAGCTTTGTTGATGCCCCCCACTTGATTTCTCTTTACCGGCTAGTTTGTGCTAATCGCTGTGTGGTGCCGTTTTTTCTCCACGAATTAGTCTGAATTTAGCCCCAATTTTGAGCTGCCATGCGTTTAATCTGCCGTAGCGATC :  598

                     *       620         *       640         *       660         *       680         *       700         *       720         *       740         *
TaDmc1-A1 : ACTGTCTGTAGAATCCTTTCGAATTCGGCACTAGATACAGTTCAATCGCGTGGGTTTTCTGTGGAAACTTGGACCAATTACGGACGCGTTTTCGTCCACTTCGTGGCGTAGGGTTT-AAGCCTGTCTTCCTCATCTGTTCCACTTTTTTT :  748
TaDmc1-B1 : ACTGGCTGTAGAATCCTGTCGAATTCGAAACTAGATACAGTTCAATCGCGTGGGTTTTCTGTGGAAGCTTGGACCAATTACGGACGCGTTTTCGTGCACTTCGAGGCGTAGGGTTTTAAGCCTGTCGTCCTCATCTGTTCCACTTTTTTT :  748
TaDmc1-D1 : ACTGGCTGTAGAATCCTTTCGAATTCGAAACTAGATACAGTTCAATCGCGTGGGTTTTCTGTGGAAACTTGGACCAATTACGGACGCGTTTTCCTGCACTTCGAGGCGTAGGGTTTTAAGCCTGTCTTCCTTATCTGTTCCACTTTTTTT :  748

                   760         *       780         *       800         *       820         *       840         *       860         *       880         *       900
TaDmc1-A1 : TTTCTGAATGTTCACCTGTTCTGTTCCATAATTACGGGTAGCTTCCGCGAGTTTAGCAAGTTTACTTTAAACATTTGTCTAGATGTGATCCGCGCTTGTGTTTTGTTTGCGTGATCCAATTGGTTCGTCGTGCACTTTGTTCATTGCCAT :  898
TaDmc1-B1 : TTTCTGAATGTTCACCTGTTCTGTTCCATAATCACGGGTAGCTTCCGCGAGTTTAGCAAGTTTACTTTAAACATTTGCCTAGATGTGATCCGCGCTTGTGTTTTGTTTGCGTGATCCAATTGGTTCGTCGTGCACTTTGTTCATTGCCAT :  898
TaDmc1-D1 : TT-CTGAATGTTCACCTGTTATGTTCCATAATTACGGGTAGCTTCCGCGAGTTTAGCAAGTTTACTTTAAACATTTGCCTAGATGTGATCCGCGCTTGTGTTTTGTTTGCGTGATCCAATTGGTTCGTCGTGCACTTTGTTCATTGCCAT :  897

                     *       920         *       940         *       960         *       980         *      1000         *      1020         *      1040         *
TaDmc1-A1 : GAATTTGAATCCTAAGCCGTAACCTTCCTTTTAACGCTGTAACTGATCACTCGTCCAAACAGTGATCTCGCAGGGAATAAACTCAGGAGACGTGAAGAAGCTGCAGGATGCGGGGATCTACACTTGCAATGGGCTGATGATGCACACCAA : 1048
TaDmc1-B1 : GAATTTGAATCCTAAGCCGT------------AACGCTGTAACTAATCACTCGTCCAAACAGTGATCTCGCAGGGAATTAACTCAGGAGACGTGAAGAAGCTGCAGGATGCGGGGATCTACACTTGCAATGGCCTGATGATGCACACCAA : 1036
TaDmc1-D1 : GAATTTGAATCCTAAGCCGTAACATTCCTTGTAACGCTGTAACTAATCACTCGTCCAAACAGTGATCTCGCAGGGAATAAACTCAGGAGACGTGAAGAAGCTGCAGGATGCGGGGATCTACACTTGCAATGGGCTGATGATGCACACCAA : 1047

                  1060         *      1080         *      1100         *      1120         *      1140         *      1160         *      1180         *      1200
TaDmc1-A1 : GAAGGTCCCAATCCCGTCAGAGCATAAATCTGCACGCATTCTCCCTAAAATTTGTGGTTGCATACTGAAGCTGATTTCTGGTGCACGACACATGACTGTTATTAGTTGCTTTATGTATTCAAGTTTCAACACATGATTGATGTTGTTTCA : 1198
TaDmc1-B1 : GAAGGTCAAACTCCCTTCAGAGCCTAAATCTGCATGCATTCTCCCTAAAATTTGTGGTTGCATACTGAAGTTGATTTCTGGTGCACGACACATGACTGTTATTAGTTGTTTTATGTATTCAAGTTTCAACACATGATTGATGTTGTTTCA : 1186
TaDmc1-D1 : GAAGGTCAAAATCCCTTCAGAGCCTAAATCTGCACGCATTCTCCCTAAAATTTGTGGTTGCATGCTGAAGTTGATTTCTGGTGCACGACACATGACTGTTATTAGTAGCTTTATGTATTCAAGTTTCAACACATGATTGATGTTGTTTCA : 1197

                     *      1220         *      1240         *      1260         *      1280         *      1300         *      1320         *      1340         *
TaDmc1-A1 : GTACCAATGTATGGATTCATCTTGCAGAGCCTTACAGGGATTAAGGGCTTGTCTGAAGCAAAGGTTGATAAGATCTGCGAGGCTGCTGAAAAACTTCTGGTATGATTGTTATCATCTTTGCATTGATTCTGGTTTACAACTTCTGTGCCA : 1348
TaDmc1-B1 : GTATCAATGTATGGATTCATCTTGCAGAGCCTTACAGGGATTAAGGGCTTGTCTGAAGCAAAGGTTGATAAGATCTGCGAGGCTGCTGAAAAACTTCTGGTATGATTGTTATCATCTTTGCATTGATTCTGGTTTAGAACTTCTGTGCCA : 1336
TaDmc1-D1 : GTATCAATGTACGGATTCATCTTGCAGAGCCTTACAGGGATTAAGGGCTTGTCTGAAGCGAAGGTTGATAAGATCTGTGAGGCTGCTGAAAAACTTCTGGTATGATTGTTATCATCTTTGCATTGATTCTGGTTTAGAACTTCTGTGCCA : 1347

                  1360         *      1380         *      1400         *      1420         *      1440         *      1460         *      1480         *      1500
TaDmc1-A1 : TATTATTCTCTTTGCATTGATTCTGGTTTACAACTTCTGTGCCATATTATTCTCTTTGCATTGATTCTGGTTTACAACTTCTGTGCCATCTGTGCTATGCGATCGAGTCTTCTGAAATTTGTATTCTATTTGATTTTCAGAGTCAGGGTT : 1498
TaDmc1-B1 : TA--------------------------------------------TTATTCTCTTTGCATTGATTCTGGTTTACAACTTCTGTGACATCTGTGCTATACGATCGAGTCTTCTGAAATTTGTATTCTATTTGATTTTCAGAGTCAGGGTT : 1442
TaDmc1-D1 : TA--------------------------------------------TTATTCTCTTTGCATTGATTCTGGTTTACAACCTCTGTGCCATCTGTGCTATGCGATAGAATCTTCTGAAGTTTGTGTACTATTTGATTTTCAGAGTCAGGGTT : 1453

                     *      1520         *      1540         *      1560         *      1580         *      1600         *      1620         *      1640         *
TaDmc1-A1 : TCATGACAGGAAGTGATCTCCTTATTAAGGTAAGGTTTAGGGAGCTAAGCTACTGATGAAGGACGACCATACAGTTTAGTGCTGTTTTTGCAGTAGTGCTACACCGCTACTATGCTAAACTTACTGTAA----TAGTTTGTTTTTGGAAT : 1644
TaDmc1-B1 : TCATGACAGGAAGTGATCTCCTTATTAAGGTAAGGTTTAGGGAGCTAAGCTACTGATGAAGGACGATCATACAGTTTAGTGCTGTTTTTGCAGTAGTGCTACACTGCTACTATGCTAAACTTACTGTTA----TAGTTTGTTTTTGGAAT : 1588
TaDmc1-D1 : TCATGACAGGAAGTGATCTCCTTATTAAGGTGAGGTTTAGG-AGCTAAGCTACTGATTAAGGACGATCATACAGTTTAGTACTGTTTTGGCCATAGTG--------CTACTATGCTAAACTTACTGTAAATAATAGTTTGTTTTTGGAAT : 1594

                  1660         *      1680         *      1700         *      1720         *      1740         *      1760         *      1780         *      1800
TaDmc1-A1 : CTGCTTGAGTGCTTTCATTACTCCACCTGTGTACTGCCCAGTTGTTTGATTTACTTTTTCGTATATTGAAGCGAAAGTCTGTTGTCCGGATTACCACTGGGAGCCAAGCGCTTGATGAGCTGCTTGGAGGTAACATATGTCGCCCTTGAT : 1794
TaDmc1-B1 : CTGCTTGAGTGCTTTCATTACTCCACCTGTGTACTGCCCGGTTGTTTGATTTACTTTTCCGTATATTGAAGCGAAAGTCTGTTGTCCGGATTACCACTGGGAGCCAAGCGCTTGATGAGCTGCTTGGAGGTAACATATGTCGTTCTTGAT : 1738
TaDMC1-D1 : CTGCTTGAGTGCTTTCATTACTCCACCTGTGTACTGCCCGGTTGTTTGATTTACTTTTCCGTATATTGAAGCGAAAGTCTGTTGTCCGGATTACCACTGGGAGCCAAACGCTTGATGAGCTGCTTGGAGGTAACATATGTCATCCTTGAT : 1744

                     *      1820         *      1840         *      1860         *      1880         *      1900         *      1920         *      1940         *
TaDmc1-A1 : TCTGTTCTGATTATTTCTGATGTTATGCTCTAACCCATTCACATATTTCCATAATTTGAAGGAGGGATTGAAACACTCTGTATCACAGAGGCATTTGGAGAGTTCCGGTCAGTAAATGTTCCAGTTACCATTTTCTTCTGGATTT----- : 1939
TaDmc1-B1 : TCTGTTCTGATTATTTCTGATGTTATGCTCTAACCTATTCACATATTTCCATAATTTGAAGGAGGGATTGAAACACTCTGTATCACAGAGGCATTTGGAGAGTTCCGGTCAGTAAATGTTCCGGTTACCATTTTCTTCTGGATTTTT--- : 1885
TaDmc1-D1 : TCTGTTCTGATTATTCCTGATGTTATGCTCTAACCTATTCAAATATTTGCGTAATTTGAAGGAGGGATTGAAACACTCTGTATCACAGAGGCATTTGGAGAGTTCCGGTGAGTAAATGTTCCCGTTACCATTTTCTTCTTGGTTTCTTCT : 1894

                  1960         *      1980         *      2000         *      2020         *      2040         *      2060         *      2080         *      2100
TaDmc1-A1 : TTTTTTGCAGGGAGCGCCTATTAGTTTTATCTCTGTTATGTATGATTTGTGCCATATTTCGCAGTAGATGTAGCAATTTTTAGTCCTAGTAATTTTTTAGT---TGGT----AAACTCTGATAAGCTGCTTTCACCTCCTTTTCTGCTGT : 2082
TaDmc1-B1 : TTTTTTGCAGGGGGCGCCTATTAGTTGTATCTCTGTTATGTATGATTTGTGCCATATTTGGCAGTAGATGTAACAATTTTTAGTCCTAGTAATTTTTTAGT---TGGT---AAAACTCTGATAAGCTTCTTTCACCTCCTTTTCTGCTGT : 2029
TaDmc1-D1 : TTTTTTGCAGGGACCGCCTATTAGTTGTAGCTCTGTTATGTATGATTTGTGCCATATTTGGCAGTAG------------CTAGTCCTAGTAATTTTTTAGTACATGGTGCTAAAACTCTCATAAGCTTCTTTCACCTCCTTTTCTACTGT : 2032

                     *      2120         *      2140         *      2160         *      2180         *      2200         *      2220         *      2240         *
TaDmc1-A1 : CTAGTCTTCCAGAGGTATTGATGATGGTGCATAT-------------------------------------------------------------------------------------------------------------------- : 2116
TaDmc1-B1 : CTAGCCTTCCAGAGGTATTGATGATAGTGCATATAACAACTCCTAAGCATGGTTGCTGTGATGCTAACATTGAATACTGGATACCTGTTACATGCCAAAATTCAAACATATCTACATCAATATTATTATTCACTATTTCTCTGATGCAAC : 2179
TaDmc1-D1 : CTAGTCTTCCAGAGGTATTGATGGTAGTGCATATAAGAATTCGAAAGCATGGTTGCTGTGATGCTAACATTGAATACTGGATACATGTTGCATGCCAAAACTCAAACAAATCTACATCAATATTACTAT-CACTATTTCCCTGATGCAAC : 2181

                  2260         *      2280         *      2300         *      2320         *      2340         *      2360         *      2380         *      2400
TaDmc1-A1 : ------------------------------------------------------------------------------------TCAGTTCCAACTTTCTAGCGCTAAAATCCATATTATTTACTTTGTCTTGCAGGTCAGGGAAGACCC : 2182
TaDmc1-B1 : ACACACATGCTCTTATTATTATTCAATGAGCACGGTTGTAATGTTGACTTCCAAAGCACATTTGTTACTCAAACATTTCTCAATTCAGTTCCAACTTTCTAGCGCTAAAATCCATATTATTTACTTTGTTTTGCAGGTCAGGGAAGACCC : 2329
TaDmc1-D1 : ACGCACCTGCTCTTATTATT-------GAGCACAGTTGTAATGTTGATTTCCAAAGCACATTTGTTACTCAACCATTTC-----TCAATTCCAACTTTCTAGCGTTAAAATCCATATTATTTACTTTGTCTTGCAGGTCAGGGAAGACCC : 2319

                     *      2420         *      2440         *      2460         *      2480         *      2500         *      2520         *      2540         *
TaDmc1-A1 : AGTTGGCTCATACTCTTTGTGTCTCCACTCAGGTCCATTTCCTGCCTTGTATATTCTCGGTGAAACCTCACTACATCAGAATCCATGAATAACTCTGCTTGTTTAATCAAA--------------------------------------- : 2293
TaDmc1-B1 : AGTTGGCTCATACTCTTTGTGTCTCCACTCAGGTCCATTTCCTGCCTTGTAT-TTCTCGGTGAAACCTCACTACATCAGAATCCATGAATAACTCTGCTTGTTTAATCAAAGAGTGAAGTGCACCCTAGGTCCTCGAACTATTTTGAAGG : 2478
TaDmc1-D1 : AGTTGGCTCATACTCTTTGTGTCTCCACTCAGGTCCATTTCCTGCCTTGTATTTTATCGATGAAACCTCACTAGAACAGAATCCATGAATAACTCTCCATGTTTAATCAAA--------------------------------------- : 2430

                  2560         *      2580         *      2600         *      2620         *      2640         *      2660         *      2680         *      2700
TaDmc1-A1 : ------------------------------------------------------------------------------------------------------------------------------------------------------ :    -
TaDmc1-B1 : TGTCATATAGGTCCTCGAATTATGAAAAGTGTCATCCAGGTCCTCAAAAATCCTTGAAGTGCAATAAGTGTGTACGTATGTGACACATTTGAAATAGTTTAAGGACCTACATGACACCGCTAAAATAGTTCGAGGACTTGGATGACACGC : 2628
TaDmc1-D1 : ------------------------------------------------------------------------------------------------------------------------------------------------------ :    -


                     *      2720         *      2740         *      2760         *      2780         *      2800         *      2820         *      2840         *
TaDmc1-A1 : ---------------------------------------------------------------------------------------------------------TGTATAACAGCTTCCACTCCACATGCATGGTGGGAACGGGAAGGT : 2338
TaDmc1-B1 : ATATTGCACTTTGAGGACCTGAATGACCCTTTTCATAGTTCGAGGACCTACGTAACACCTTTTAAATAGTTTGAGGACCTTTGATGCACTTCACTCTTAATCAAATGTATAACAGCTTCCACTCCATATGCATGGTGGGAACGGGAAGGT : 2778
TaDmc1-D1 : ---------------------------------------------------------------------------------------------------------TGTATAACAGCTTCCACTCCACATGCATGGTGGGAACGGGAAGGT : 2475
                                                                                                                     TGTATAACAGCTTCCACTCCAcATGCATGGTGGGAACGGGAAGGT

                  2860         *      2880         *      2900         *      2920         *      2940         *      2960         *      2980         *      3000
TaDmc1-A1 : TGCCTACATTGACACTGAGGGAACATTGTATCCTTTGAATTCCTTAGTAATACCTATAGTCGATTTGTTCAATGAATTCATTTACATTCCTGTGCTTATCGAAACTATTCCCTTAAAGGAGTTATCAGCCGGCCTGAACGCATTGTGCCA : 2488
TaDmc1-B1 : TGCCTACATTGGCACTGAGGGAACGTTGTATCCTTTGAATTCCTTAGTAATACCTAT-----------TTAATGAATTCATTTACATTCCTGTATTTATCGAAACTGTTCCCTTAACGGAGTTATCAGCCGGCCTGAACGCATTGTGCCA : 2917
TaDmc1-D1 : TGCCTACATTGACACTGAGGGAACATTGTATCCTTTGAATTCCTTAGTAATACCTATAGCTGATTTGTTCGATGAATTCATTTACATTCCTGTATTTCTCGAAACTGTTCCCTTAACGGAGTTATCAGCCGGCCTGAACGCATTGTGCCA : 2625

                     *      3020         *      3040         *      3060         *      3080         *      3100         *      3120         *      3140         *
TaDmc1-A1 : ATTGCTGAGAGATTTGGGATGGATGCCAATGCTGTTCTTGACAATGTATGGCTCCTTTTACATCTCTCTTA--ACCCATTTAAGGAAGAATAGATCAAGATCTTTGTTTAATTCGTGATCTTTCTGTTTTAGATCATATACGCTCGCGCA : 2636
TaDmc1-B1 : ATTGCTGAGAGATTTGGGATGGATGCCAATGCTGTTCTTGACAATGCATGGCTCCTTTTACATCTCTCTTA--ACCCATTTAAGGAAGAATAGATAAAGATCTTTGTTTAATTCGTGATCTTTCTGTTTTAGATCATATACGCTCGTGCA : 3065
TaDmc1-D1 : ATTGCTGAGAGATTTGGGATGGATGCCAATGCTGTTCTTGACAATGTATGGGTCCTTTTACATCTCTCTTATAACCCATTTAAGGAAGAATAGATCGAGATCTTTGTTTAATTTGTGATCTTTCTGTTTTAGATCATATATGCTCGTGCC : 2775

                  3160         *      3180         *      3200         *      3220         *      3240         *      3260         *      3280         *      3300
TaDmc1-A1 : TACACCTATGAGCACCAGTACAACTTACTCCTGGGCCTTGCTGCCAAGATGGCCGAAGAGCCTTTCAGGCTTCTGGTACGCATGACTTTGCTGACATGTACTATCAAACTTACAAGTTGATAGATCTCAACTGTGCTCATGTGATCTTTG : 2786
TaDmc1-B1 : TACACCTATGAGCACCAGTACAACTTACTCCTGGGCCTTGTTGCCAAGATGGCTGAAGAGCCTTTCAGGCTTCTGGTACGCATGACTTTGCTGACATGTAGTGTTAAACTTACAAGTTGATAGATCTCAACTGTGCTCATGTAATCTTTG : 3215
TaDmc1-D1 : TACACCTATGAGCACCAGTACAACTTACTCCTGGGCCTTGCTGCCAAGATGGCTGAAGAGCCTTTCAGGCTTCTGGTACGCATGACTTTGCTGCCATGT------AAATTTACAAGTTGATAGATCTCAACTGTGCTCATGTGTTCTTTG : 2919

                     *      3320         *      3340         *      3360         *      3380         *      3400         *      3420         *      3440         *
TaDmc1-A1 : TTTGGCTTGGAAATGATAGATCGTGGATTCTGTGATTGCGCTATTCCGTGTTGATTTCAGTGGTAGGGGTGAACTTGCAGAGCGTCAGGTATTCTACTGTAACTAGCTAACTATGTGAAAAAATCAAGCAACTCATGATGTAGTCGAATG : 2936
TaDmc1-B1 : TTTGGCTTGGAAATGATAGATCGTGGATTCTGTGATTGCGCTATTCCGTGTTGATTTCAGTGGCAGGGGTGAACTTGCAGAGCGTCAGGTATTCTACTGTAACTAGCTAACTATGTGAAAAAATCAAGCAACTCATGATGTAGTCGAATG : 3365
TaDmc1-D1 : TTTGACTTGGAAATGATAGATCGTGGATTCTGTGATTGCGCTGTTCCGTGTTGATTTCAGTGGTAGGGGTGAACTTGCAGAGCGTCAGGTATTCTACTGTAACTAGCTAACTACATGAAAAAATCAAGCAACTCATGAAGTAGTCGAATG : 3069

                  3460         *      3480         *      3500         *      3520         *      3540         *      3560         *      3580         *      3600
TaDmc1-A1 : CT---TGCATTTTATACACTTGCTCTAAGTGATGTGCTCTGGAACTGCAGCAAAAACTGGCACAAATGCTGTCCCGCCTTACAAAGATTGCTGAGGAGTTCAATGTTGCAGTGTACATCACCAACCAAGGTGTGCTTT-CCAATCTATCC : 3082
TaDmc1-B1 : CT---TGCATTTTATACACTTGCTCTAAGTGATGTGCTCTGGAACTGCAGCAAAAACTGGCACAAATGCTTTCCCGCCTTACAAAGATTGCTGAGGAGTTCAATGTTGCAGTGTACATCACCAACCAAGGTGTGCTTTTCCAATCTATCC : 3512
TaDmc1-D1 : CTGCTTGCATTTTATACACTTGCTTTAAGTGATGTGCTCTGGAACTGCAGCAAAAGCTGGCACAAATGCTGTCCCGCCTTACAAAGATTGCTGAGGAGTTCAATGTTGCAGTGTACATCACCAACCAAGGTGTGCTTT-CCAATCTATCC : 3218

                     *      3620         *      3640         *      3660         *      3680         *      3700         *      3720         *      3740         *
TaDmc1-A1 : TGTCTGTTCCAAGAAAGAGCTCCTTATATTCGTGGATCTCAAATCATATAAGTTCTTTCCTTGTTCCAGTGATTGCGGACCCAGGTGGTGGTATGTTCATCACTGACCCCAAAAAGCCGGCAGGAGGCCACGTGCTGGCGCATGCAGCCA : 3232
TaDmc1-B1 : TGTCTGTTCCAAGAAAGAGCTCCTTATATTCGTGGATCTCAAATCATACAAGTTCTTTCCTTGTTCCAGTGATTGCGGACCCAGGTGGTGGTATGTTCATCACTGACCCCAAAAAGCCGGCGGGAGGCCACGTGCTGGCGCATGCAGCCA : 3662
TaDmc1-D1 : TGTCTGTTCCAAGAAAGAGCTCCTTATATTCGTGGATCTCAAATCATATAAGTTCTTTCCTTGTTCCAGTGATTGCGGACCCAGGTGGTGGTATGTTCATCACTGACCCCAAAAAGCCGGCGGGAGGCCACGTGCTGGCGCATGCAGCCA : 3368


                  3760         *      3780         *      3800         *      3820         *      3840         *      3860         *      3880         *      3900
TaDmc1-A1 : CCATCCGGTTGATGCTGAGGAAAAGCAAAGGCGAGCAGCGTGTCTGCAAGATCTTTGACGCCCCTAACCTTCCCGAGGGAGAAGCTATATCCTTTTGCT---TACTA------CTTGTTTACTGCTTGTGC--TATTCATTGCTTTAATT : 3371
TaDmc1-B1 : CCATCCGGTTGATGCTGAGGAAAGGCAAAGGCGAGCAGCGTATCTGCAAGATCTTTGACGCCCCTAACCTTCCCGAGGGAGAAGCTATATCCTTTTGCTCTATACTAGGAGTACTTGTTTACTGCTTGTGC--TATTCATTCCATTAATT : 3810
TaDmc1-D1 : CCATCCGGTTGATGCTGAGGAAAGGCAAAGGCGAGCAGCGTGTCTGCAAGATCTTTGACGCCCCTAACCTTCCCGAGGGAGAAGCTATATCCTTTTGCTCTATACTA------CTTGTTTACTGCTTGTGCGCTATTCATTGCTTTAATT : 3512

                     *      3920         *      3940         *      3960         *      3980         *      4000         *      4020         *      4040         *
TaDmc1-A1 : TGTTTGGTTGCTGAACTCTTGATAGGATTGTTTTGCTCAGACTTGCACAGC----TTTATGATTTAGTTCTTAGTTGTTAACCTCCTTTGCTGAAGCTTCACGTAGCACAAATTAGCAATGATCAGATGTCGAGAAATTGTTTCCCTTGA : 3517
TaDmc1-B1 : TGTTTGGTTGCTGAACTCTTGATAGGATTGTTTTGCTCAGACTTGCGCAGCAAAGTTTATGATTTAGTTCT-----------------------------------------------------AGTTGTTGAGAAATTGTTTCGCTTGA : 3907
TaDmc1-D1 : TGTTTGGTTGTTGAACTCTTGATAGGATTGTTTTGCTCAGACTTGCACAGCAAAGTTTATGATTTAGTTCT-AGTTGTTAACCTCCTTTACTGAAGCTTCACGTAGCACAAATTAGCGACGATCAGATGTCGAGAAATTGTTTCCCTTGA : 3661

                  4060         *      4080         *      4100
TaDmc1-A1 : CCACCAACACGTTTTCCAGATTACAACAGGCGGATTGATGGATGTGAAAGACTGA : 3572
TaDmc1-B1 : CCACCAACACGTTTTCCAGATTACAACAGGTGGATTGATGGATGTGAAAGACTGA : 3962
TaDmc1-D1 : CCACCAACACGTTTTCCAGATCACAACAGGCGGATTGATGGATGTGAAAGACTGA : 3716

```
