## Supplemental Fig. 4 for "*Dmc1* is a candidate for temperature tolerance during wheat meiosis"

### Slide 1
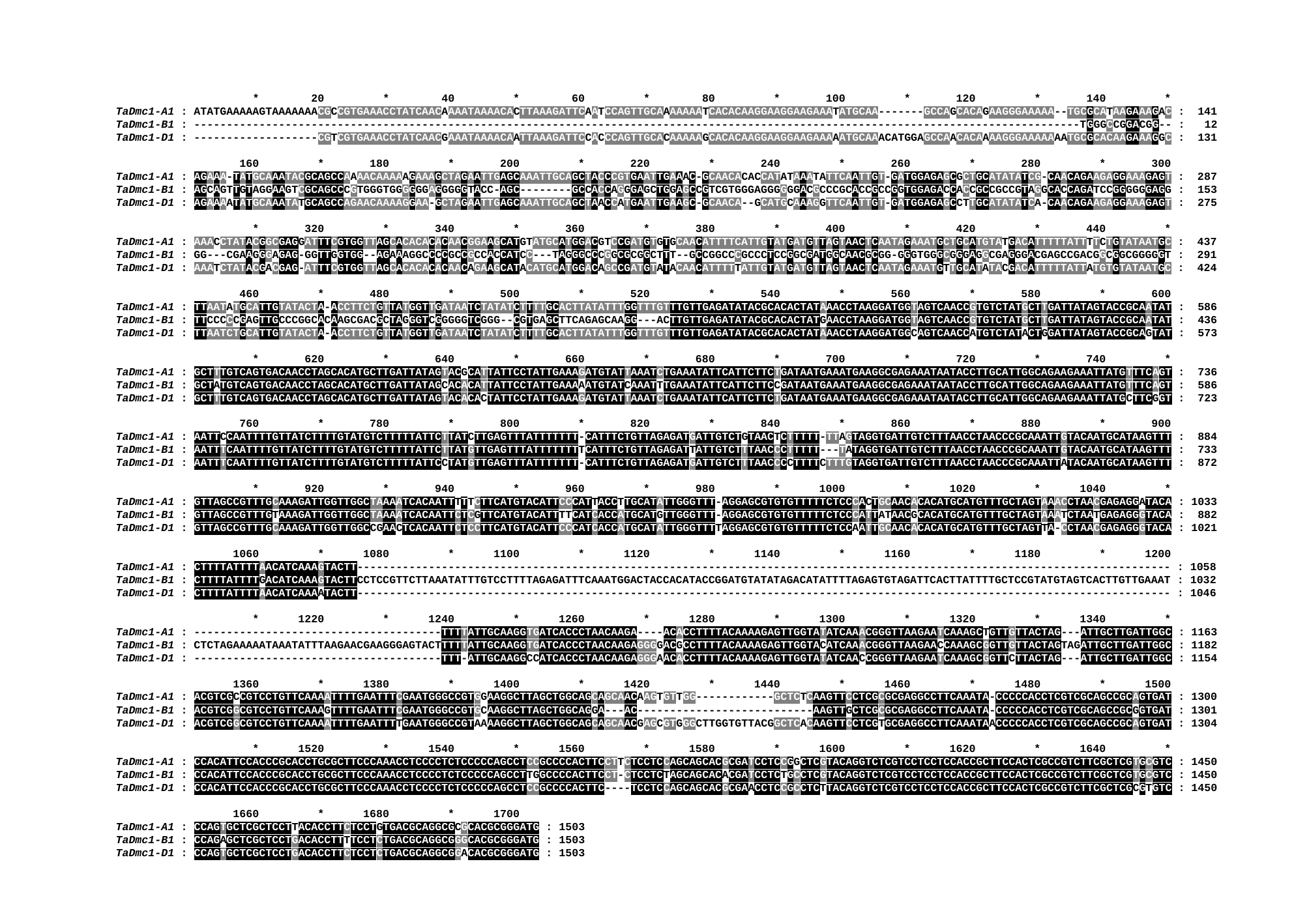

* 20 * 40 * 60 * 80 * 100 * 120 * 140 *
TaDmc1-A1 : ATATGAAAAAGTAAAAAAACGCCGTGAAACCTATCAACAAAATAAAACACTTAAAGATTCAATCCAGTTGCAAAAAAATCACACAAGGAAGGAAGAAATATGCAA-------GCCAGCACAGAAGGGAAAAA--TGCGCATAAGAAAGAC : 141
TaDmc1-B1 : ----------------------------------------------------------------------------------------------------------------------------------------TGGGCCGGACGG-- : 12
TaDmc1-D1 : -------------------CGTCGTGAAACCTATCAACGAAATAAAACAATTAAAGATTCCACCCAGTTGCACAAAAAGCACACAAGGAAGGAAGAAAAATGCAAACATGGAGCCAACACAAAAGGGAAAAAAATGCGCACAAGAAAGGC : 131
 160 * 180 * 200 * 220 * 240 * 260 * 280 * 300
TaDmc1-A1 : AGAAA-TATGCAAATACGCAGCCAAAACAAAAAGAAAGCTAGAATTGAGCAAATTGCAGCTACCCGTGAATTGAAAC-GCAACACACCATATAAATATTCAATTGT-GATGGAGAGCGCTGCATATATCG-CAACAGAAGAGGAAAGAGT : 287
TaDmc1-B1 : AGCAGTTGTAGGAAGTCGCAGCCCGTGGGTGGGGGGAGGGGGTACC-AGC--------GCCACCAGGGAGCTGGAGCCGTCGTGGGAGGGGGGACGCCCGCACCGCCGGTGGAGACCACCGCCGCCGTAGGCACCAGATCCGGGGGGAGG : 153
TaDmc1-D1 : AGAAAATATGCAAATATGCAGCCAGAACAAAAGGAA-GCTAGAATTGAGCAAATTGCAGCTAACCATGAATTGAAGC-GCAACA--GCATGCAAAGGTTCAATTGT-GATGGAGAGCCTTGCATATATCA-CAACAGAAGAGGAAAGAGT : 275
 * 320 * 340 * 360 * 380 * 400 * 420 * 440 *
TaDmc1-A1 : AAACCTATACGGCGAGGATTTCGTGGTTAGCACACACACAACGGAAGCATGTATGCATGGACGTCCGATGTGTGCAACATTTTCATTGTATGATGTTAGTAACTCAATAGAAATGCTGCATGTATGACATTTTTATTTTCTGTATAATGC : 437
TaDmc1-B1 : GG---CGAAGGGAGAG-GGTTGGTGG--AGAAAGGCCCCGCCGCCACCATCC---TAGGGCCCGGCGCGGCTTT--GCCGGCCCGCCCTCCGGCGATGGCAACGCGG-GGGTGGGCGGGAGGCGAGGGACGAGCCGACGGCGGCGGGGGT : 291
TaDmc1-D1 : AAATCTATACGACGAG-ATTTCGTGGTTAGCACACACACAACAGAAGCATACATGCATGGACAGCCGATGTATACAACATTTTTATTGTATGATGTTAGTAACTCAATAGAAATGTTGCATATACGACATTTTTATTATGTGTATAATGC : 424
 460 * 480 * 500 * 520 * 540 * 560 * 580 * 600
TaDmc1-A1 : TTAATATGCATTGTATACTA-ACCTTCTGTTATGGTTGATAATCTATATCTTTTGCACTTATATTTGGTTTGTTTGTTGAGATATACGCACACTATAAACCTAAGGATGGTAGTCAACCGTGTCTATGCTTGATTATAGTACCGCAATAT : 586
TaDmc1-B1 : TTCCCCCGAGTTGCCCGGCACAAGCGACGCTAGGGTCGGGGGTCGGG--CGTGAGCTTCAGAGCAAGG---ACTTGTTGAGATATACGCACACTATGAACCTAAGGATGGTAGTCAACCGTGTCTATGCTTGATTATAGTACCGCAATAT : 436
TaDmc1-D1 : TTAATCTGCATTGTATACTA-ACCTTCTGTTATGGTTGATAATCTATATCTTTTGCACTTATATTTGGTTTGTTTGTTGAGATATACGCACACTATAAACCTAAGGATGGCAGTCAACCATGTCTATACTGGATTATAGTACCGCAGTAT : 573
 * 620 * 640 * 660 * 680 * 700 * 720 * 740 *
TaDmc1-A1 : GCTTTGTCAGTGACAACCTAGCACATGCTTGATTATAGTACGCATTATTCCTATTGAAAGATGTATTAAATCTGAAATATTCATTCTTCTGATAATGAAATGAAGGCGAGAAATAATACCTTGCATTGGCAGAAGAAATTATGTTTCAGT : 736
TaDmc1-B1 : GCTATGTCAGTGACAACCTAGCACATGCTTGATTATAGCACACATTATTCCTATTGAAAAATGTATCAAATTTGAAATATTCATTCTTCCGATAATGAAATGAAGGCGAGAAATAATACCTTGCATTGGCAGAAGAAATTATGTTTCAGT : 586
TaDmc1-D1 : GCTTTGTCAGTGACAACCTAGCACATGCTTGATTATAGTACACACTATTCCTATTGAAAGATGTATTAAATCTGAAATATTCATTCTTCTGATAATGAAATGAAGGCGAGAAATAATACCTTGCATTGGCAGAAGAAATTATGCTTCGGT : 723
 760 * 780 * 800 * 820 * 840 * 860 * 880 * 900
TaDmc1-A1 : AATTCCAATTTTGTTATCTTTTGTATGTCTTTTTATTCTTATCTTGAGTTTATTTTTTT-CATTTCTGTTAGAGATGATTGTCTGTAACTCTTTTT-TTAGTAGGTGATTGTCTTTAACCTAACCCGCAAATTGTACAATGCATAAGTTT : 884
TaDmc1-B1 : AATTTCAATTTTGTTATCTTTTGTATGTCTTTTTATTCTTATGTTGAGTTTATTTTTTTTCATTTCTGTTAGAGATTATTGTCTTTAACCCTTTTT---TATAGGTGATTGTCTTTAACCTAACCCGCAAATTGTACAATGCATAAGTTT : 733
TaDmc1-D1 : AATTTCAATTTTGTTATCTTTTGTATGTCTTTTTATTCCTATGTTGAGTTTATTTTTTT-CATTTCTGTTAGAGATGATTGTCTTTAACCCCTTTTCTTTGTAGGTGATTGTCTTTAACCTAACCCGCAAATTATACAATGCATAAGTTT : 872
 * 920 * 940 * 960 * 980 * 1000 * 1020 * 1040 *
TaDmc1-A1 : GTTAGCCGTTTGCAAAGATTGGTTGGCTAAAATCACAATTTTTCTTCATGTACATTCCCATTACCTTGCATATTGGGTTT-AGGAGCGTGTGTTTTTCTCCCACTGCAACACACATGCATGTTTGCTAGTAAACCTAACGAGAGGATACA : 1033
TaDmc1-B1 : GTTAGCCGTTTGTAAAGATTGGTTGGCTAAAATCACAATTCTCGTTCATGTACATTTTCATCACCATGCATGTTGGGTTT-AGGAGCGTGTGTTTTTCTCCCATTATAACGCACATGCATGTTTGCTAGTAAATCTAATGAGAGGGTACA : 882
TaDmc1-D1 : GTTAGCCGTTTGCAAAGATTGGTTGGCCGAACTCACAATTCTCCTTCATGTACATTCCCATCACCATGCATATTGGGTTTTAGGAGCGTGTGTTTTTCTCCAATTGCAACACACATGCATGTTTGCTAGTTA-CCTAACGAGAGGGTACA : 1021
 1060 * 1080 * 1100 * 1120 * 1140 * 1160 * 1180 * 1200
TaDmc1-A1 : CTTTTATTTTAACATCAAAGTACTT----------------------------------------------------------------------------------------------------------------------------- : 1058
TaDmc1-B1 : CTTTTATTTTGACATCAAAGTACTTCCTCCGTTCTTAAATATTTGTCCTTTTAGAGATTTCAAATGGACTACCACATACCGGATGTATATAGACATATTTTAGAGTGTAGATTCACTTATTTTGCTCCGTATGTAGTCACTTGTTGAAAT : 1032
TaDmc1-D1 : CTTTTATTTTAACATCAAAATACTT----------------------------------------------------------------------------------------------------------------------------- : 1046
 * 1220 * 1240 * 1260 * 1280 * 1300 * 1320 * 1340 *
TaDmc1-A1 : --------------------------------------TTTTATTGCAAGGTGATCACCCTAACAAGA----ACACCTTTTACAAAAGAGTTGGTATATCAAACGGGTTAAGAATCAAAGCTGTTGTTACTAG---ATTGCTTGATTGGC : 1163
TaDmc1-B1 : CTCTAGAAAAATAAATATTTAAGAACGAAGGGAGTACTTTTTATTGCAAGGTGATCACCCTAACAAGAGGGGACGCCTTTTACAAAAGAGTTGGTACATCAAACGGGTTAAGAACCAAAGCGGTTGTTACTAGTAGATTGCTTGATTGGC : 1182
TaDmc1-D1 : --------------------------------------TTT-ATTGCAAGGCCATCACCCTAACAAGAGGGAACACCTTTTACAAAAGAGTTGGTATATCAACCGGGTTAAGAATCAAAGCGGTTCTTACTAG---ATTGCTTGATTGGC : 1154
 1360 * 1380 * 1400 * 1420 * 1440 * 1460 * 1480 * 1500
TaDmc1-A1 : ACGTCGCCGTCCTGTTCAAAATTTTGAATTTCGAATGGGCCGTGGAAGGCTTAGCTGGCAGCAGCAACAAGTGTTGG------------GCTCTCAAGTTCCTCGCGCGAGGCCTTCAAATA-CCCCCACCTCGTCGCAGCCGCAGTGAT : 1300
TaDmc1-B1 : ACGTCGGCGTCCTGTTCAAAGTTTTGAATTTCGAATGGGCCGTGCAAGGCTTAGCTGGCAGGA---AC---------------------------AAGTTGCTCGCGCGAGGCCTTCAAATA-CCCCCACCTCGTCGCAGCCGCGGTGAT : 1301
TaDmc1-D1 : ACGTCGGCGTCCTGTTCAAAATTTTGAATTTTGAATGGGCCGTAAAAGGCTTAGCTGGCAGCAGCAACGAGCGTGGGCTTGGTGTTACGGCTCACAAGTTCCTCGTGCGAGGCCTTCAAATAACCCCCACCTCGTCGCAGCCGCAGTGAT : 1304
 * 1520 * 1540 * 1560 * 1580 * 1600 * 1620 * 1640 *
TaDmc1-A1 : CCACATTCCACCCGCACCTGCGCTTCCCAAACCTCCCCTCTCCCCCAGCCTCCGCCCCACTTCCTTCTCCTCCAGCAGCACGCGATCCTCCGGCTCGTACAGGTCTCGTCCTCCTCCACCGCTTCCACTCGCCGTCTTCGCTCGTGCGTC : 1450
TaDmc1-B1 : CCACATTCCACCCGCACCTGCGCTTCCCAAACCTCCCCTCTCCCCCAGCCTTGGCCCCACTTCCT-CTCCTCTAGCAGCACACGATCCTCTGCCTCGTACAGGTCTCGTCCTCCTCCACCGCTTCCACTCGCCGTCTTCGCTCGTGCGTC : 1450
TaDmc1-D1 : CCACATTCCACCCGCACCTGCGCTTCCCAAACCTCCCCTCTCCCCCAGCCTCCGCCCCACTTC----TCCTCCAGCAGCACGCGAACCTCCGCCTCTTACAGGTCTCGTCCTCCTCCACCGCTTCCACTCGCCGTCTTCGCTCGCGTGTC : 1450
 1660 * 1680 * 1700
TaDmc1-A1 : CCAGTGCTCGCTCCTTACACCTTCTCCTGTGACGCAGGCGCGCACGCGGGATG : 1503
TaDmc1-B1 : CCAGAGCTCGCTCCTGACACCTTTTCCTCTGACGCAGGCGGGCACGCGGGATG : 1503
TaDmc1-D1 : CCAGTGCTCGCTCCTGACACCTTCTCCTCTGACGCAGGCGGACACGCGGGATG : 1503
